# Erlin1/2 Complex is a Dynamic Scaffold for Membrane Protein Sequestration and Microdomain Assembly on the Endoplasmic Reticulum

**DOI:** 10.1101/2025.06.14.659634

**Authors:** Lu Yan, Zihong Xu, Yuanhang Yao, Xiaoting Wang, Yonglun Wang, Chengying Ma, Ningning Li, Xiaowei Chen, Ning Gao

**Affiliations:** State Key Laboratory of Membrane Biology, Peking-Tsinghua Joint Center for Life Sciences, Academy for Advanced Interdisciplinary Studies, School of Life Sciences, Peking University, Beijing 100871, China; State Key Laboratory of Membrane Biology and Institute of Molecular Medicine, College of Future Technology, Peking University, Beijing 100871, China; National Biomedical Imaging Center, Peking University, Beijing 100871, China; Changping Laboratory, Beijing 102206, China; Beijing Advanced Center of RNA Biology (BEACON), Peking University, Beijing 100871, China

**Author notes:** These authors contributed equally (Lu Yan, Zihong Xu, Yuanhang Yao). Correspondence (X.C.); (N.G.).

**Keywords:** Cryo-electron microscopy (cryo-EM), SPFH, erlin1/2, Functional membrane microdomain (FMM), endoplasmic reticulum-associated degradation (ERAD), β-coronavirus

## Abstract

The SPFH (Stomatin, Prohibitin, Flotillin, and HflK/C) family of proteins are key scaffolding components involved in the organization of functional membrane microdomains (FMMs) across various subcellular membranes, including those of the endoplasmic reticulum (ER) and mitochondria, which are characterized by a low content of saturated lipids. Among this protein family, the erlin1/2 complex is specifically located on the ER membrane. Previous studies have shown that the erlin1/2 complex plays an essential role in the ER-associated degradation (ERAD) pathway, mediating the ubiquitin-dependent degradation of various proteins such as inositol 1,4,5-trisphosphate receptors (InsP3Rs), which are important calcium ion transporters on the ER membrane. However, the molecular mechanisms underlying erlin-mediated FMMs organization and its role in ERAD remain poorly understood. In this study, we determined the single-particle cryo-electron microscopy (cryo-EM) structure of the erlin1/2 complex under different detergent conditions. Our findings reveal that the erlin1/2 complex forms a 26-mer cage-like structure, composed of alternating erlin1 and erlin2 subunits. The erlin1/2 complex defines a nanodomain on the ER membrane, which could recruit various types of proteins to both the interior and exterior membrane regions of the cage. By caging cargo proteins, the erlin1/2 complex physically secludes them from their binding partners, leading to a potential halt in their function. Moreover, individual cages can further interact with one another, facilitating the organization of FMMs of different sizes on the ER membrane. These dynamic properties may play a general and critical role in various processes occurring on the ER, including viral replication, positioning the erlin1/2 complex as a promising new target for antiviral drug development.

## Introduction

Lipids and proteins in cell membranes exhibit lateral heterogeneity, forming dynamic, nanometer-sized microdomains enriched with saturated lipids such as cholesterol and glycosphingolipids^1–4^. These regions, known as detergent-resistant membranes (DRMs), are characterized by their low solubility in non-ionic detergents at low temperatures. Various proteins involved in signal transduction, immune response, and pathogen invasion have been found to be enriched in DRMs, suggesting that these functional membrane microdomains (FMMs) serve as an important organizational unit in biological membranes^2,5–8^. However, due to their highly dynamic nature and heterogeneous composition, the organization and regulatory mechanisms of FMMs remain largely unknown.

The SPFH (Stomatin, Prohibitin, Flotillin, and HflK/C) protein family is widely distributed across eukaryotes and prokaryotes, and is found in the plasma membrane (PM) and various subcellular membranes, including lipid droplets, endosomes, lysosomes, mitochondria and the endoplasmic reticulum (ER), which is characterized by a low contect of saturated lipids^9–11^. Members of this family have been detected in DRMs and are thought to function as scaffolding proteins for FMMs^10^. In mammals, the SPFH family primarily includes stomatin^12^, stomatin-like proteins^13^, prohibitin1/2^14^, flotillin1/2^15^, podocin^16^, and erlin1/2^11^, which are the only members that reside in the ER membrane^11^. The identification of erlin1 and erlin2 has led to the hypothesis that FMMs with high lipid order may also exist within the ER^17^.

The erlin1/2 complex has been shown to play key roles in the ER-associated degradation (ERAD) pathway, a central protein quality control system within the ER^18,19^. The ERAD pathway not only targets misfolded proteins for degradation to maintain ER homeostasis, but also responds to physiological signals, such as cholesterol levels in the ER membrane, by degrading key enzymes involved in lipid biosynthesis^20–22^. This contributes to the regulation of lipid metabolism. Previous studies have demonstrated that the erlin1/2 complex can rapidly associate with activated inositol 1,4,5-trisphosphate receptors (InsP3Rs), serving as scaffolds that bridge InsP3Rs and E3 ubiquitin ligases like RNF170, thereby facilitating their ubiquitination and subsequent degradation^23–27^. The erlin1/2 complex also participates in the ERAD-associated degradation of 3-hydroxy-3-methylglutaryl-CoA (HMG-CoA) reductase, a key enzyme in cholesterol synthesis^28^. Furthermore, the erlin1/2 complex inhibits the ERAD-mediated degradation of Insig1 under cholesterol-replete conditions, thereby retaining the SCAP-SREBP complex on the ER membrane and limiting the activation of SREBPs to maintain lipid homeostasis^20,29^. These findings highlight the role of the erlin1/2 complex in regulating lipid homeostasis within the ER. Additionally, mutations in the erlin1/2 complex have been linked to neurodegenerative disorders^30–33^.

Erlin1 and erlin2 share high sequence similarity and are thought to assemble into high-molecular-weight, ring-like complexes. However, evidence also suggest that erlin1 and erlin2 monomers are capable of forming homo-oligomeric assemblies^18,34^. Despite these insights, structural evidence elucidating the molecular mechanisms underlying their functional roles remains insufficient. It is still unclear whether the erlin1/2 complex requires both proteins to form a functional heterologous complex or whether erlin1 and erlin2 can exhibit independent stability.

In this study, we found that neither erlin1 nor erlin2 alone is sufficient to form a complete cage-like homo-oligomeric complex. Instead, erlin1 and erlin2 assemble into a 26-mer cage-like structure in an alternating arrangement. Under mild non-ionic detergent conditions, we observed that the cage structure could contract, potentially altering the curvature of the membrane confined within it. Furthermore, various cargo proteins were found to bind to both the interior and exterior surfaces of the erlin1/2 cages, and closely packed pairs of erlin1/2 particles were observed, suggesting that the erlin1/2 complex may act as an organizational hub for different ER-associated proteins. Our findings provide structural evidence that the erlin1/2 complex serves as a scaffold for FMMs on the ER membrane, offering new insights for further studies of ER FMMs in processes such as lipid synthesis, viral replication, vesicle transport, and other biological functions.

## Results

### Erlin1 and erlin2 co-assemble to form a closed cage complex

Erlin1 and erlin2 are two homologous proteins with ~72% sequence identity and extremely high (~90%) sequence similarity (Figure S1). Previous studies have indicated that the depletion of erlin1 or erlin2 leads to slightly different phenotypes^18^, and they exhibit independent stability^34^. Unlike flotillin1/2^35^ and prohibitin1/2^36^, it has also been proposed that erlin1 and erlin2 may be capable of forming both homo- and hetero-complexes to fulfill their respective functions. To investigate this, we overexpressed and purified erlin1 and erlin2 proteins separately, each fused with a C-terminal Flag tag, from human HEK293F cells. The membrane proteins were solubilized using a 3:1 (w/w) mixture of DDM and GDN, and the proteins were then eluted into pure GDN. The enriched samples were further separated by a 10%–50% glycerol density gradient to isolate potential erlin1 or erlin2 complexes.

Surprisingly, despite the high sequence similarity between erlin1 and erlin2 (Figure S1), their particles exhibited distinct sizes, as determined by 10%–50% glycerol gradient centrifugation (Figure 1A), and distinct morphologies, as observed by negative-staining electron microscopy (nsEM) (Figures 1B and 1C). Erlin1 was enriched at approximately 35% glycerol concentration (Figure 1A), and its particles predominantly displayed an arc shape with a diameter of around 40 nm (Figure 1B). In contrast, erlin2 oligomers had a lower molecular weight and peaked at around 25% glycerol concentration in the gradient (Figure 1A), with particles showing a linear structure with a length of approximately 80 nm (Figure 1C).

**Figure 1.**
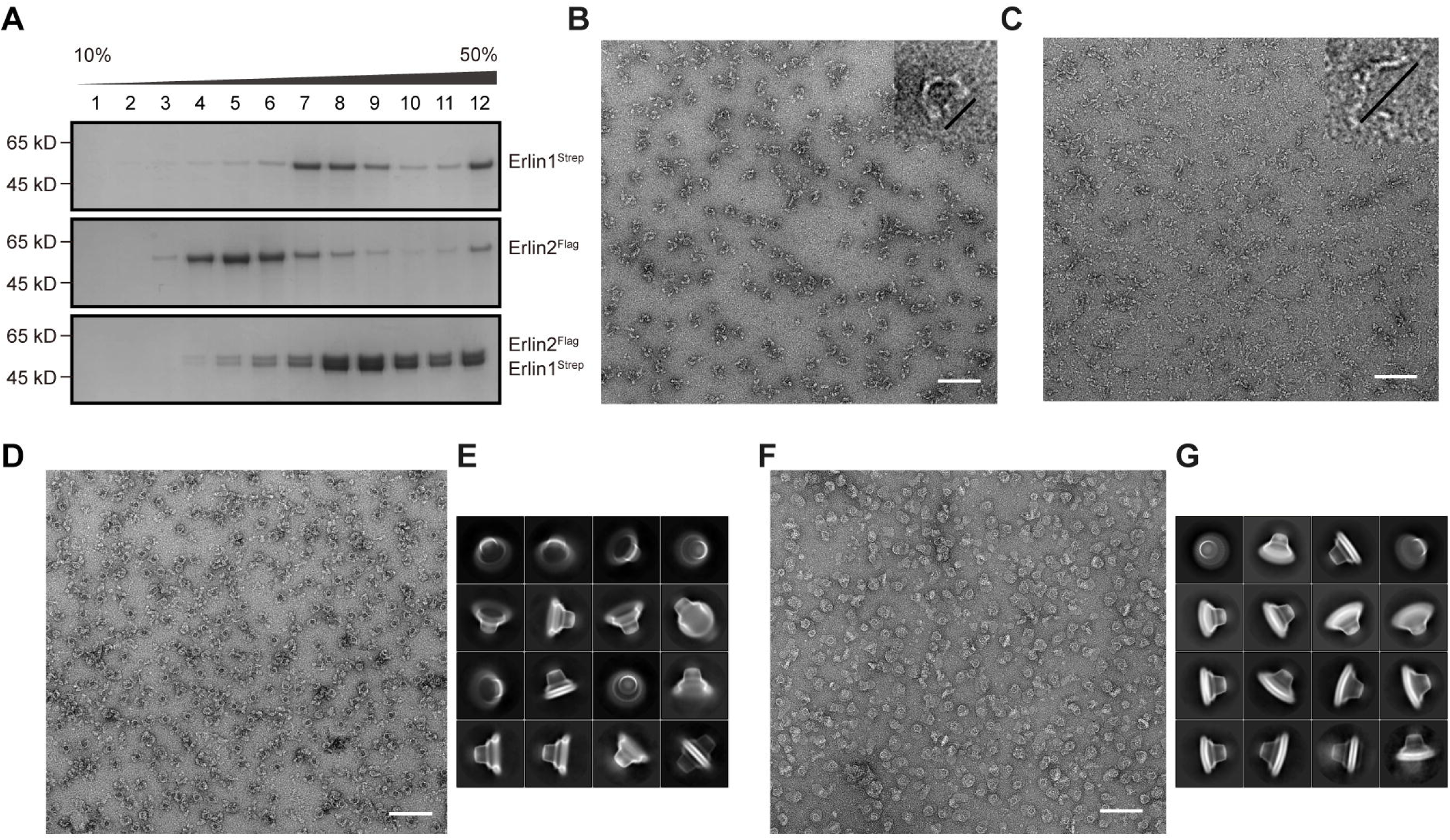
Purification and structural characterization of erlin1, erlin2 and the erlin1/2 complexes. (A) Coomassie Brilliant Blue-stained 10%–50% glycerol gradient fractions of purified proteins from individually overexpressed erlin1, erlin2, and co-overexpressed erlin1 and erlin2. Erlin1 alone forms higher-order oligomers enriched in fractions 7–9, while erlin2 forms lower-order oligomers enriched in fractions 4–6. The erlin1/2 complexes also appear as higher-order oligomers, predominantly distributed in fractions 8–10. (B and C) Negative-staining EM (nsEM) images of individually overexpressed erlin1 and erlin2. Erlin1 forms a structure composed of multiple arc-shaped subunits with a diameter of approximately 40 nm (B, upper-right inset). Erlin2 displays a linear conformation with a length of approximately 80 nm (C, upper-right inset). Scale bar: 200 nm. (D) nsEM images of erlin1/2 complexes purified under the DDM+GDN solubilization condition. Scale bar: 200 nm. (E) Cryo-EM 2D classification of erlin1/2 complexes in (D). The complex adopts a closed, cage-like architecture, most of the particles contain no detectable membrane components inside the cage. A minor population of particles exhibits a double-cage organization. (F) nsEM images of erlin1/2 complexes purified under the GDN+CHS solubilization condition. Scale bar: 200 nm. (G) Cryo-EM 2D classification of erlin1/2 complexes in (F). The complex also adopts a complete cage-like structure but retains membrane components within the cage. Particles also display both contracted and expanded conformational states.

Given that several members of the SPFH family, such as HflK/C in *E. coli*^37,38^ and flotillin1/2 in the human plasma membrane^39^, require both subunits to form cage-like structures, we hypothesized that erlin1/2 may do the same. To test this, we co-overexpressed erlin1 and erlin2, with a Flag tag fused to erlin2 and a twin-Strep tag fused to erlin1, connected by a P2A sequence for co-translational expression. Both Strep- and Flag-based affinity purifications successfully enriched the erlin1/2 complex (Figures S2A–S2C). The complexes were high-molecular-weight oligomers, enriched at approximately 40% concentration in the glycerol density gradient (Figure 1A), and the nsEM analysis revealed that they formed a relatively complete circular structure (Figure 1D).

We also tested various detergent conditions for membrane protein purification (Figures 1D–1G and S2D–S2F). Although DDM improved protein yield, it negatively impacted the integrity of the erlin1/2 complex (Figures S2E and S2F). Therefore, we chose two purification conditions. The first used a mixture of DDM and GDN (3:1, w/w) for solubilization, which maximized protein yield, and pure GDN for elution (hereafter referred to as DDM+GDN for simplicity). The second used a mixture of GDN and CHS (10:1, w/w) to better preserve the membrane components associated with the cage complexes (hereafter referred to as GDN+CHS for simplicity).

We then collected and processed single-particle cryo-EM data for these samples purified by both methods. From the 2D classification results, we observed that membrane components bound to the complex were largely displaced when DDM was used during membrane solubilization (Figures 1D and 1E). In contrast, the double-layered membrane components in the complexes were well-preserved with the GDN+CHS method, and the membranes exhibited variability in size and curvature (Figures 1F and 1G). Furthermore, 2D and 3D classification revealed that the cage-like structures prepared with GDN+CHS exhibited slightly different diameters; among them, the structures of the expanded and contracted cages were solved at resolutions of 6.11 Å and 2.76 Å, respectively (Figure S3 and S5). Particles from the DDM+GDN sample were more uniform, and the final structure was solved at a resolution of 2.60 Å (Figure S4 and S5).

### Erlin1 and erlin2 form a heterogeneous 26-mer cage-like structure through alternating assembly

We compared the structure purified with DDM+GDN with the two states obtained with GDN+CHS and found that, while the overall sizes of the three cages differed slightly, the stoichiometry of the complexes remained the same, forming a heterogeneous 26-mer cage-like structure. The membrane spanning region of the cage has a diameter of approximately 24 nm and an extramembrane height of around 15 nm (Figures 2A and 2B). Notably, the three structures could be consistently aligned in the C-terminal region (Figure S6). Similar to HflK/C^37,38^,each subunit of erlin1/2 is composed of four domains: the N-terminal transmembrane domain (TM), the SPFH domain (SPFH1 and SPFH2), the coiled-coil domain (CC1 and CC2), and the C-terminal domain (Figure 2C). The TM and SPFH1 domains are responsible for membrane insertion, the SPFH2 domain forms the cage base, the long CC1 domain constitutes the cage wall, and the short CC2 and the C-terminal domain form the cage top.

**Figure 2.**
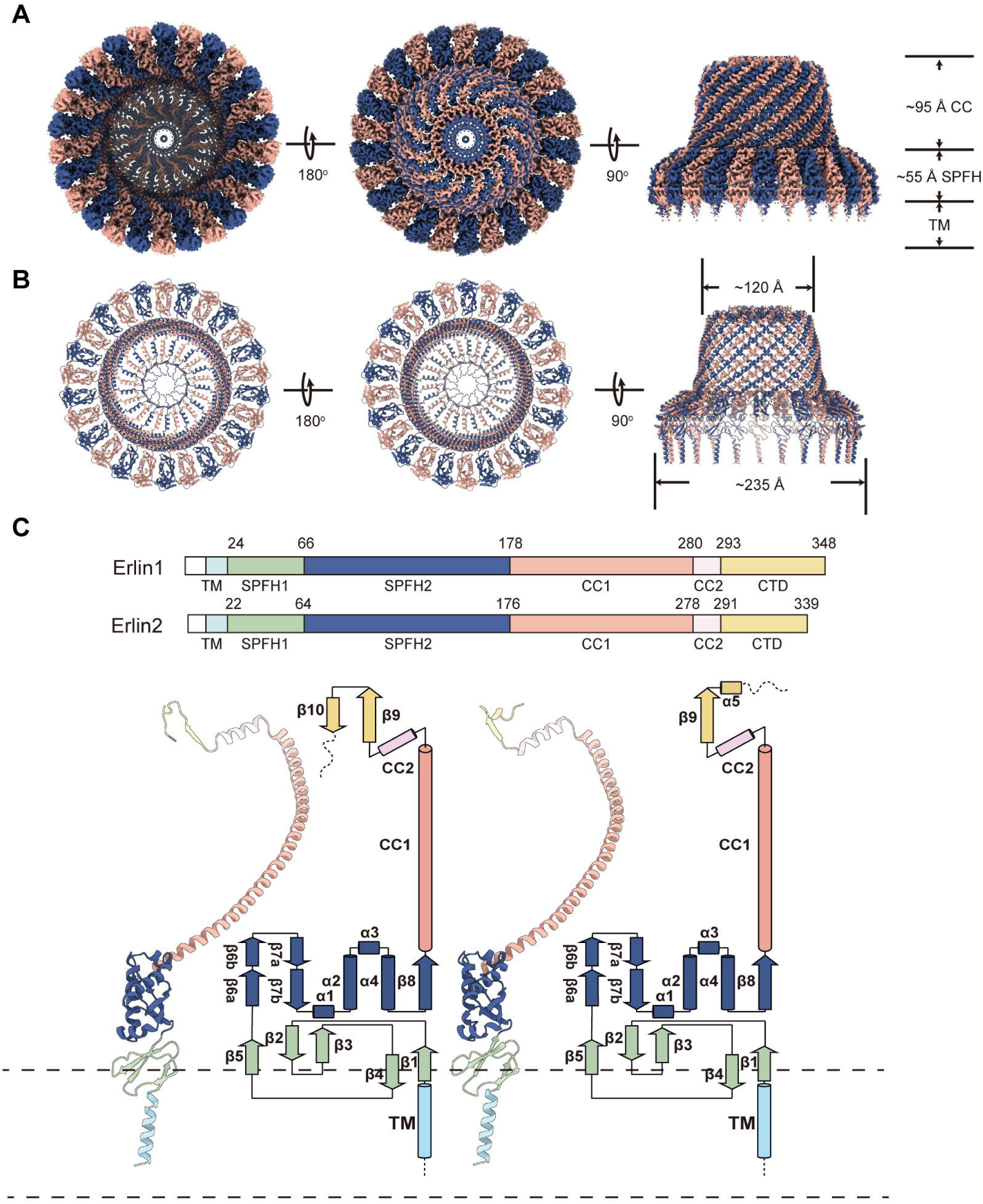
Overall structure of the erlin1/2 complex. (A) Bottom (left), top (middle), and side (right) views of the cryo-EM density map of the erlin1/2 complex. The cage-like structure consists of 26 subunits. The SPFH domains (SPFH1 and SPFH2) span approximately 35 Å in height, while the coiled-coil (CC) region extends approximately 72 Å. Erlin1 and erlin2 subunits are colored in blue and pink, respectively. (B) Bottom (left), top (middle), and side (right) views of the atomic model of the erlin1/2 complex. The diameters of the cage assembly are approximately 235 Å and 120 Å in the membrane-proximal and -distal regions, respectively. Erlin1 and erlin2 subunits are colored in blue and pink, respectively. (C) Schematic representation of the domain organization of erlin1 and erlin2 (top), and corresponding atomic model with topological diagram (bottom). TM, transmembrane domain, sky blue; SPFH1, SPFH1 domain, green; SPFH2, SPFH2 domain, dark blue; CC1, Coiled-coil domain 1, orange; CC2, Coiled-coil domain 2, pink; CTD, C-terminal domain, yellow.

During data processing, when refining the erlin1/2 complex as a whole, the densities of the cage top, in particular the C-termini of erlin1/2, were not sufficiently resolved for atomic modelling. Therefore, we refined the cage top separately, significantly improving its resolution (Figures S3C and S4C). From the improved maps, we observed substantial differences in the C-terminal domains of adjacent subunits. One C-terminal tail extended inward into the hydrophobic channel formed by β9-strand, while the other bent outward to form a short helical structure (Figures 3A, S7A and S7B). Based on AlphaFold predictions^40^, the C-terminus of erlin2 exactly contains a short α-helix, whereas the C-terminus of erlin1 forms a loop, confirming the structural observation that erlin1 and erlin2 assemble in a 13:13 stoichiometric complex. This assignment was also confirmed by subtle differences in the cryo-EM density map between erlin1 and erlin2 at the end of the CC1 domain (Y264 in erlin1 and C262 in erlin2) (Figure S7C). In addition, the density map revealed that both N108 of erlin1 and N106 of erlin2 were glycosylated (Figure S7D), confirming their localization in the ER lumen, and each subunit contains an intramolecular disulfide bond, C78-C130 and C76-C128 for erlin1 and erlin2, respectively.

**Figure 3.**
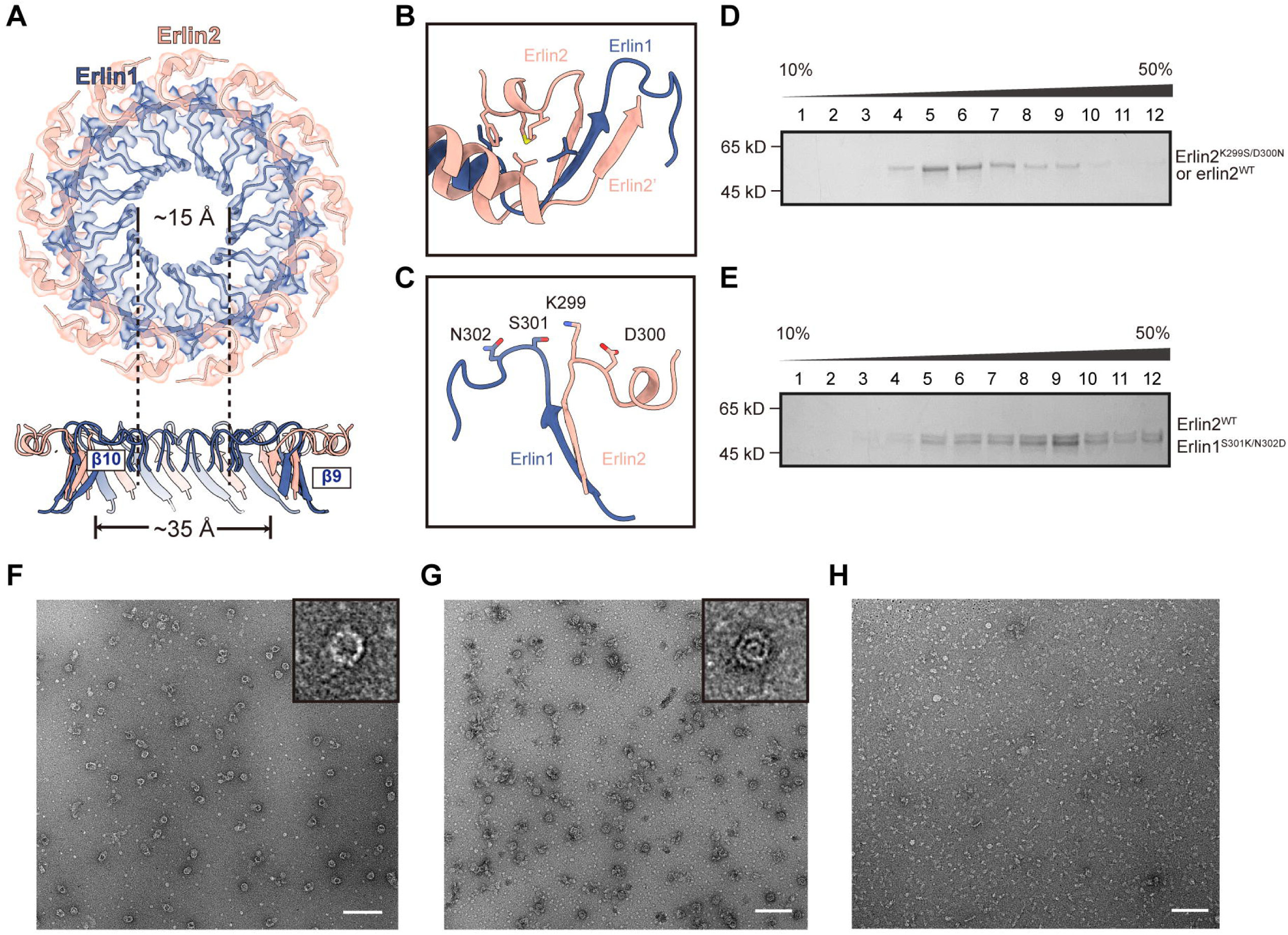
Topological organization of the erlin1/2 cage in the membrane-distal region. (A) Top and side views of the cap region of the erlin1/2 complex. At the top of the cage, after the outer β9-barrel with a diameter of approximately 35 Å, erlin2 (orange) bends outward while erlin1 (blue) bends inward to form the innermost β10-barrel wall of the central channel of the cage top with a diameter of approximately 15 Å. (B) A short C-terminal α-helix of erlin2 is stabilized by hydrophobic interactions with the CC2 region. (C) Residues located at the C-terminal turn after the β9-strand, which are not conserved between erlin1 and erlin2 but are important for complex assembly are shown. (D) Coomassie Brilliant Blue-stained 10%–50% glycerol gradient fractions of purified erlin2^WT^-erlin2^K299S/D300N^ complexes, most of which exist in a lower-order oligomeric state, with a minor fraction forming higher-order complexes at fraction 9. (E) Coomassie Brilliant Blue-stained 10%–50% glycerol gradient fractions of purified erlin1^S301K/N302D^-erlin2^WT^ complexes, most of which still form higher-order assemblies. (F–H) nsEM analysis of mutant erlin1/2 complexes. Particles from the higher-order fractions (fraction 9) of the erlin2^WT^-erlin2^K299S/D300N^ sample form cage-like structures (F). For the erlin2^WT^-erlin1^S301K/N302D^ sample, particles adopt “Swiss roll”-like structures in the higher-order fraction (fraction 9) (G) and small linear structures in the lower-order fraction (fraction 5) (H). Scale bar: 200 nm.

Consequently, we obtained a complete atomic model for the 26-mer erlin1/2 complex. In this model, 13 erlin1 and 13 erlin2 subunits, through extensive interactions from the N-terminal SPFH1 domain all the way to the C-terminal tail, alternate to form this membrane-attached cage (Figure S8). After CC2, the β9-strands from erlin1 and erlin2 form a 26-stranded β9-barrel (Figure 3A). The C-terminus of erlin1 bends inward after the β9-strand, forming the innermost 13-stranded β10-barrel, leaving a central channel on the cage top with a diameter of approximately 15 Å (Figure 3A). In contrast, the C-terminus of erlin2 contains a short α-helix, which extends outward and binds to the CC2 domain through hydrophobic interactions, constituting the upper layer of the cage top (Figure 3B).

### The C-terminal tails of erlin1 and erlin2 play an important role in the assembly of the 26-mer complex

We also observed that approximately 20% of the erlin1/2 cages were broken (Figures S3 and S4). In these structures, the walls and bases of the cages were partially disrupted, but the C-terminal top remained intact. We speculated that the C-terminal domain plays a critical role in the complex polymerization. In agreement with this idea, previous experiments have shown that truncation of C-terminal residues of erlin2 impaired erlin1/2 complex assembly^18^. Given the observation that the C-termini of erlin1/2 exhibit sharp differences in both sequence and conformation (Figures 2 and S1), we hypothesize that they might play a role in determining the stoichiometry of the erlin1/2 complex. Correlating with our nsEM data, erlin1 alone displayed arc-shaped structures, resembling features of incomplete cages. Given that the inner and outer β-barrel layers in the cage top contains precisely 13 and 26 strands, respectively, the C-terminal tail of erlin1, when polymerizing alone, would result in a mismatch between the required subunit stoichiometry in the two β-barrel layers, preventing the formation of a closed ring structure (Figure 1B). In contrast, the particles of erlin2 alone appear smaller and thinner, likely forming lower-order oligomers. We speculated that two erlin2 monomers next to each other may experience steric conflict at the C-terminus due to positioning of the last α-helix, hindering further oligomerization. Upon analyzing the C-terminal sequences, we found some critical differences between erlin1 and erlin2 after the β9-strand, which determines the orientation of their C-termini. While erlin1 has a weakly charged S301, erlin2 contains a positively charged K299 at the equivalent position (Figure 3C). Also, N302 in erlin1 is replaced by a negatively charged residue D300 in erlin2. To test whether these residues play a role in the cage assembly, we mutated K299 and D300 of erlin2 to Ser and Asn respectively (referred to as erlin2^K299S/D300N^), corresponding to their equivalent sites in erlin1. Overexpression of erlin2^K299S/D300N^ and erlin2^WT^ proteins in equal proportions in HEK293F cells showed that although the majority of particles were lower-order oligomers, there was also a weak peak in the high molecular-weight fractions (Figure 3D). The nsEM data confirmed these higher-order particles exhibited closed circular structures similar to the erlin1/2 complex (Figure 3F), indicating that mutating these two residues altered the oligomeric state.

Next, we introduced reciprocal mutations in erlin1 (S301K/N302D), mimicking the equivalent residues in erlin2. After purifying the overexpressed erlin1^S301K/N302D^-erlin2^WT^ samples, we found that although most proteins still polymerized (Figure 3E), the top view of these resulting complexes resembled a “Swiss roll” (Figure 3G), rather than the typical ring structure seen in the WT erlin1/2 complex (Figure 1D). Additionally, a small peak appeared in fraction 5 (Figure 3E), and nsEM data showed that these particles were linear, similar to the structure formed by erlin2 alone (Figure 3H).

These results indicate that proper polymerization of the C-terminal domain is essential for the formation of the erlin1/2 complex, and for a complete 26-mer cage-like complex to form, erlin1 and erlin2 must alternate.

### The erlin1/2 complex forms a membrane microdomain on the ER membrane

In addition to the N-terminal transmembrane helix, the SPFH1 domain also contributes to the membrane anchoring of the erlin1/2 complex (Figure 4), resembling the mechanisms observed in the bacterial HflK/C complex^37,38^ and the human flotillin1/2 complex^39^. Structural analysis revealed that approximately one-third of the SPFH1 domain is embedded within the membrane (Figures 4A and 4B), with the conserved positively charged residues positioned at the membrane interface (Figure 4C). Notably, additional density was observed within the membrane insertion region of the SPFH1 domain (Figures 4A, 4B, 4D and 4E). Although the identity of this density remains unresolved, among potential candidates, the density most closely resembles the head group of phosphatidylinositol (PI), which contains a bulky myo-inositol ring (Figure 4E).

**Figure 4.**
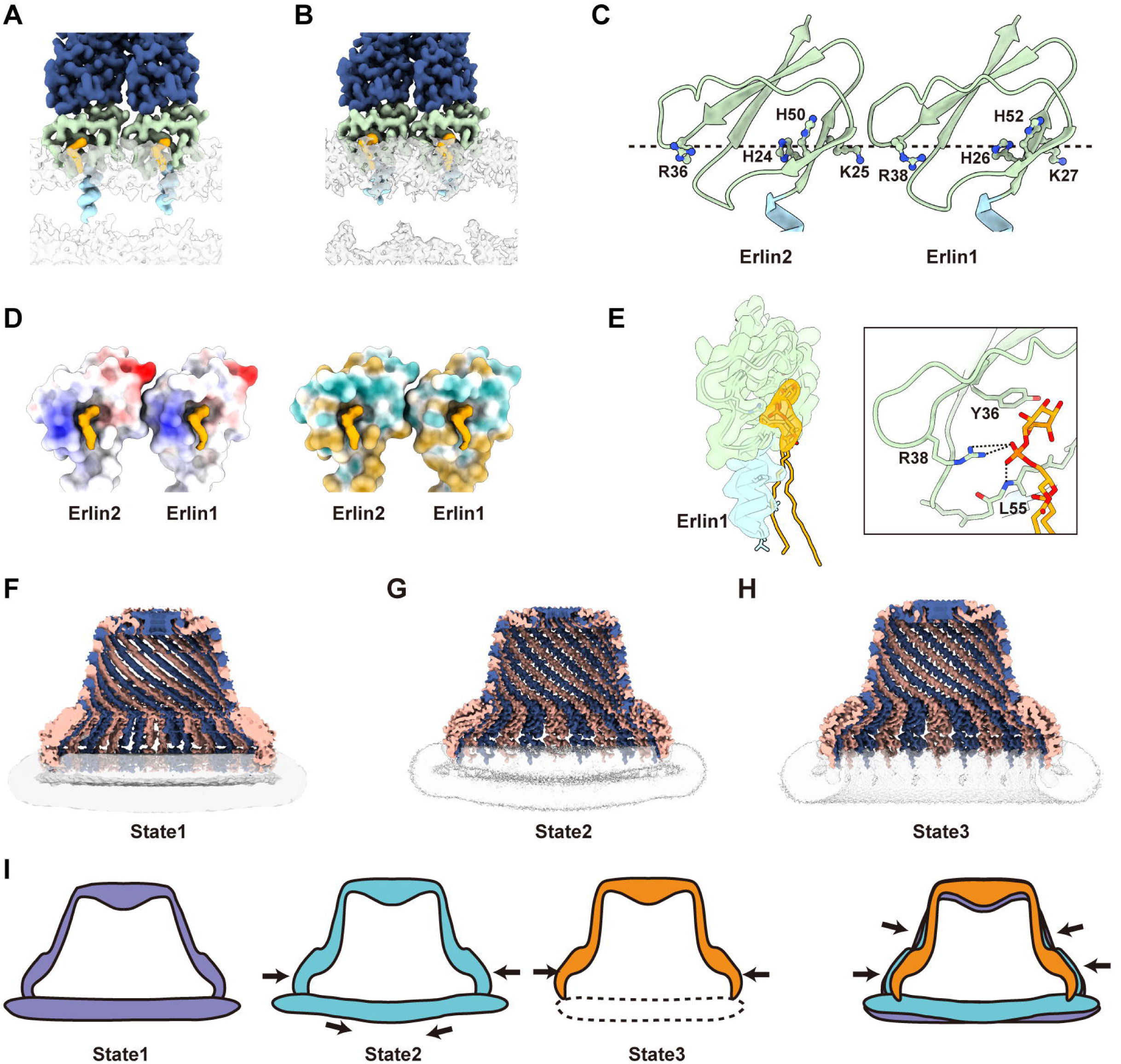
Intramembrane domains of the erlin1/2 complex. (A and B) Zoomed-in view of the TM and SPFH1 regions of the erlin1/2 complex purified by GDN+CHS (A) or DDM+GDN (B). The unidentified density embedded on the exterior surface of the SPFH1 domain is highlighted in orange. (C) Display of the membrane-inserted residues of the SPFH1 domains of the erlin1/2 complex. The conserved, positively charged residues located at the membrane interface are labeled. (D) Electrostatic (left) and hydrophobicity (right) surface representations of the SPFH1 domains of erlin1/2. The unidentified density inserts into a pocket formed by the SPFH1 domain and is positioned near a conserved, positively charged arginine residue. (E) Putative fit of a PI molecule into the unidentified density. On the basis of this fitting, the conserved residue R38 of erlin1 interacts with the phosphate group of PI, while another conserved residue Y36—unique to the erlin1/2 subfamily among all SPFH members—interacts with the *myo*-inositol ring. (F–I) Longitudinal sectional view of the cryo-EM density maps (F–H) and schematic diagrams (I) of the erlin1/2 cages in three different conformational states. State1: expanded cage (GDN+CHS); State2: contracted cage (GDN+CHS); State3: fully contracted cage after membrane dissolution (DDM+GDN).

The SPFH1 domain of erlin1/2 forms a pocket that accommodates the lipid head group (Figure 4D). Within this pocket, the phenolic hydroxyl group of a conserved tyrosine (erlin1-Y36 and erlin2-Y34)—a residue uniquely present in all erlins among SPFH family proteins—is positioned to form a stacking interaction with the inositol moiety. More importantly, the conserved arginine residues (erlin1-R38 and erlin2-R36) and main-chain N-atoms of leucines (erlin1-L55 and erlin2-L53) are precisely positioned to form electrostatic interactions with the phosphate group of the lipid, further stabilizing its binding.

Further, the SPFH1 domains of the erlin1/2 complex are embedded exclusively in the luminal leaflet of the ER membrane. In contrast, the TM domains span both leaflets, and the 26 TM helices are spaced apart and show no interaction (Figures 4F–4I). Thus, the SPFH1 domains act as physical barrier in the luminal leaflet between the inner and outer membrane regions of the erlin1/2 cage. Since the SPFH1 domains interact with only one leaflet of the membrane, this may also create a composition asymmetry between the two phospholipid layers of the ER membrane.

Comparing the three erlin1/2 structures obtained under different purification conditions, we observe that the cage-like structure can undergo contraction (Figures 4F–4I). While the C-terminal top regions of the three structures remain relatively constant, structural deflection begins to appear at the bottom of the CC1 domains and gradually increases towards the N-terminal region. On the SPFH1 domains, the difference can be as large as 9 Å (Figure S6D), indicating that the erlin1/2 cage structure can contract. The complex purified under mild detergent conditions (GDN+CHS) preserves membrane regions confined within the erlin1/2 cage, and the membrane curvature varies depending on the state of cage contraction. In the expanded state (state1 in Figures 4F and 4I), the membrane remains relatively flat, whereas in the contracted state (state2 in Figures 4G and 4I), a pronounced curvature was observed. When DDM+GDN were used to fully dissolve the membrane region enclosed within the cage, the cage could further shrink to reach its most compact state (state3 in Figures 4H and 4I). Given that non-ionic detergents are less effective at solubilizing membrane regions rich in saturated lipids, we hypothesize that the lipid components retained within the erlin1/2 cages may exhibit a high degree of order. The tightly packed, highly ordered lipids might impede the sliding of the TM helices, leading to the observed membrane curvatures. Therefore, we speculate that the erlin1/2 cage-like structure may dynamically expand and contract, in response to certain stimuli on the ER to change the local membrane curvature.

### The erlin1/2 complex binds to a variety of cargo proteins and self-organizes into clusters

Very interestingly, through examination of raw cryo-EM particles, we observed that many erlin1/2 cages exhibited obvious association with unknown proteins, varying in shapes and binding positions. Some proteins were seen to bind to the membrane region within the cage, with large extra-membrane density protruding outside the cytosolic leaflet; some were found to be trapped inside the cage; and some were seen to bind next to the cage in the outside membrane region (Figures 5A and S9A). Consistently, in the cryo-EM maps obtained using the GDN+CHS condition, the densities of transmembrane proteins were clearly visible in the membrane region inside the erlin1/2 cage (Figure S9B). Combined with the results of Coomassie Brilliant Blue staining during purification, we conclude that the cargo proteins bound to the erlin1/2 complex are highly diverse. This contrasts with the bacterial HflK/C complex, which only specifically interacts with membrane-bound AAA+ protease FtsH^37,38^.

**Figure 5.**
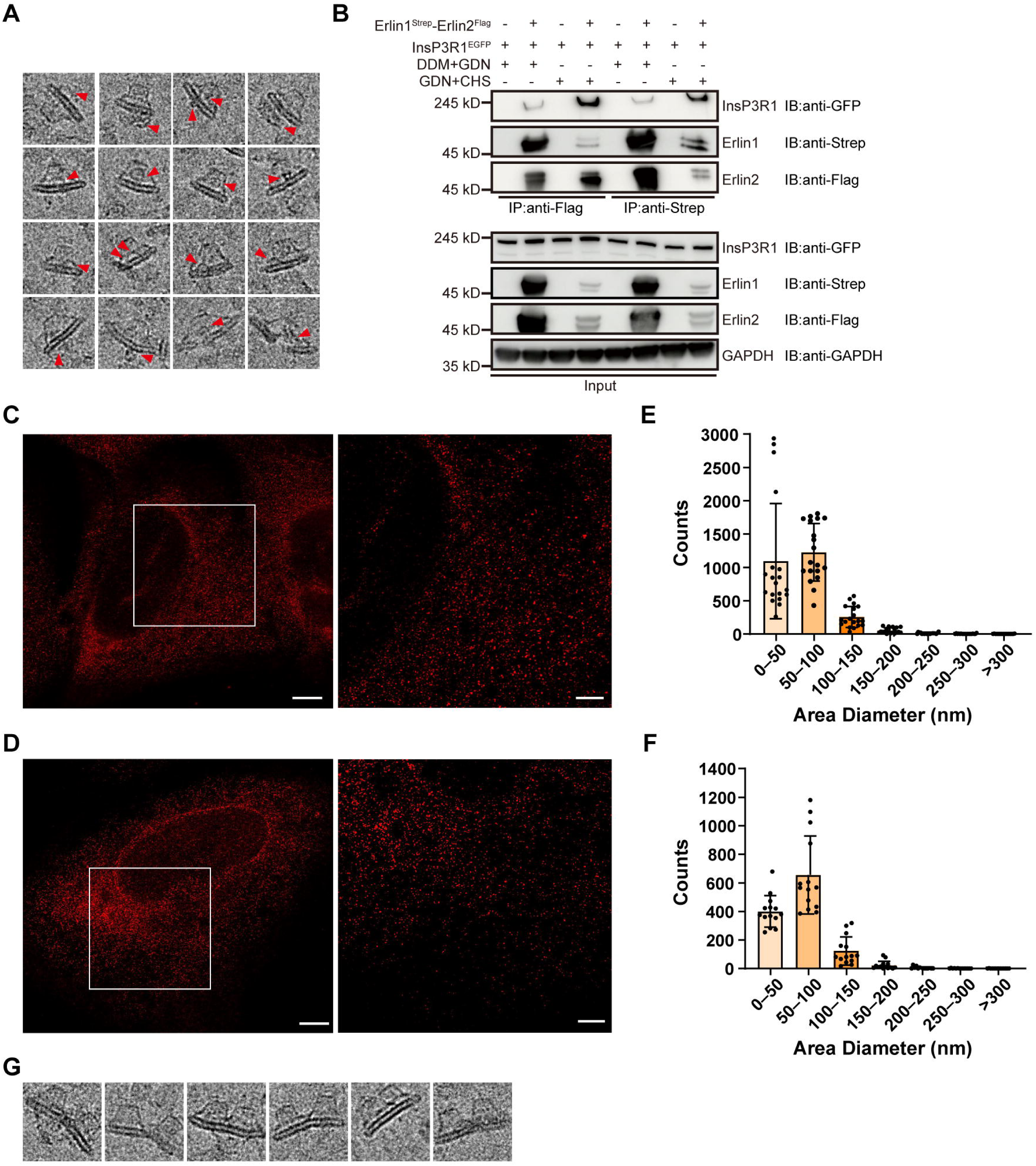
Erlin1/2-mediated membrane proteins sequestration and FMMs organization on the ER. (A) Representative raw cryo-EM particles of erlin1/2 complexes purified by GDN+CHS. Additional densities are observed either inside (the first and second rows) or adjacent (the third row) to the erlin1/2 cage. Some cages appear to be broken, with large deformations on the cage wall (the fourth row). (B) Co-expression of erlin1^Strep^-P2A-erlin2^Flag^ with InsP3R1^EGFP^ in HEK293F cells, followed by membrane lysis under varying detergent conditions—GDN+DDM or GDN+CHS—to assess interactions of erlin1/2 with cargo proteins. Co-IP was performed using affinity tags on erlin1 (Strep) or erlin2 (Flag) to evaluate their association with InsP3R1 under different solubilization conditions. Equal amounts of cells were used for these experiments. Cell lysates were subjected to membrane solubilization, and GAPDH, InsP3R1, erlin1 and erlin2 in the resulting soluble fractions were detected using respective tag antibodies (input). Immunoprecipitations were done using either anti-Flag or anti-Strep beads. (C and D) Subcellular localization of endogenous erlin1 (C) or erlin2 (D) in U2-OS cells, visualized by STED microscopy. Both proteins exhibit a punctate distribution pattern. Scale bar of whole cell image (left): 5 μm. Scale bar of zoomed-in view (right): 2 μm. (E and F) Puncta diameters in (C) and (D) were quantified using FIJI software. Statistical analysis was based on 19 (E) or 14 (F) regions from different cells. The resolution limit of the STED microscopy used was approximately 50 nm. (G) Representative raw cryo-EM particles showing clustered erlin1/2 cages.

Previous studies have demonstrated that the erlin1/2 complex plays a critical role in the ERAD pathway, particularly in mediating the ubiquitination-dependent degradation of activated InsP3Rs^18,19,25,28,29^. To further elucidate the functional contribution of erlin1/2 in this process, we selected InsP3R1 as a model cargo for mechanistic investigation. Co-expression of erlin1/2 with InsP3R1 in HEK293F cells, followed by co-immunoprecipitation (co-IP) using tagged erlin1 or erlin2, revealed differential detergent sensitivity in their interactions. Firstly, as expected, introduction of DDM, which is a better membrane solubilization agent compared with CHS, greatly enhanced the solubility of erlin1/2 complexes (Figure 5B) and consistent with previous studies, both erlin1 and erlin2 could effectively co-immunoprecipitate InsP3R1^18,19,23,26,27^. However, under the DDM+GDN condition, which disrupts membrane components associated with the erlin1/2 cage and dramatically reduces the association between erlin1/2 and InsP3R1 (Figure 5B), indicating that their interaction is more dependent on the integrity of the erlin1/2-mediated membrane microdomain and InsP3R1 is likely trapped inside the erlin1/2 cage.

Immunofluorescence imaging of endogenous erlin1 or erlin2 by stimulated emission depletion (STED) microscopy revealed their localization to discrete punctate structures on the ER membrane, mostly ~100□nm in diameter, with occasional larger puncta exceeding 300□nm (Figures 5C–5F). Additionally, we also observed in the raw cryo-EM images that some erlin1/2 particles were next to each other, forming a higher-order assembly (Figure 5G). This suggests that the cages may cluster together, potentially organizing larger FMMs on the ER membrane. Altogether, these observations suggest that the erlin1/2 complex is highly likely a general scaffold protein for organizing FMMs on the ER membrane.

### The erlin1/2 complex functions beyond ERAD and contributes to viral replication

Our observation that the erlin1/2 complex could bind various proteins on the ER membrane suggests that it may function beyond ERAD and be involves in various ER processes associated with FMMs. It is known that the FMMs on the ER are also crucial for the formation of viral replication compartments that harbor lipids and proteins necessary for viral replication^41,42^. For example, positive-strand RNA (+RNA) viruses, such as β-coronavirus (β-CoV), induce double-membrane vesicles (DMVs) tethered to the ER by thin membrane connectors to facilitate RNA replication^43^. Non-structural protein 6 (NSP6) is essential for this connection and may act as an organizer of DMV clusters^44^. To explore this further, we overexpressed the NSP6 protein of SARS-CoV-2 using adeno-associated virus (AAV) in A549 cells. Consistent with previous results^44^, confocal imaging showed that NSP6 induced the formation of the characteristic whorl-like membrane bodies on the ER, approximately 1 µm in diameter (Figure 6A). Immunostaining of endogenous erlin1 showed that it was also enriched in these NSP6-containing structures on the ER, suggesting a potential role for erlin1 in viral replication.

**Figure 6.**
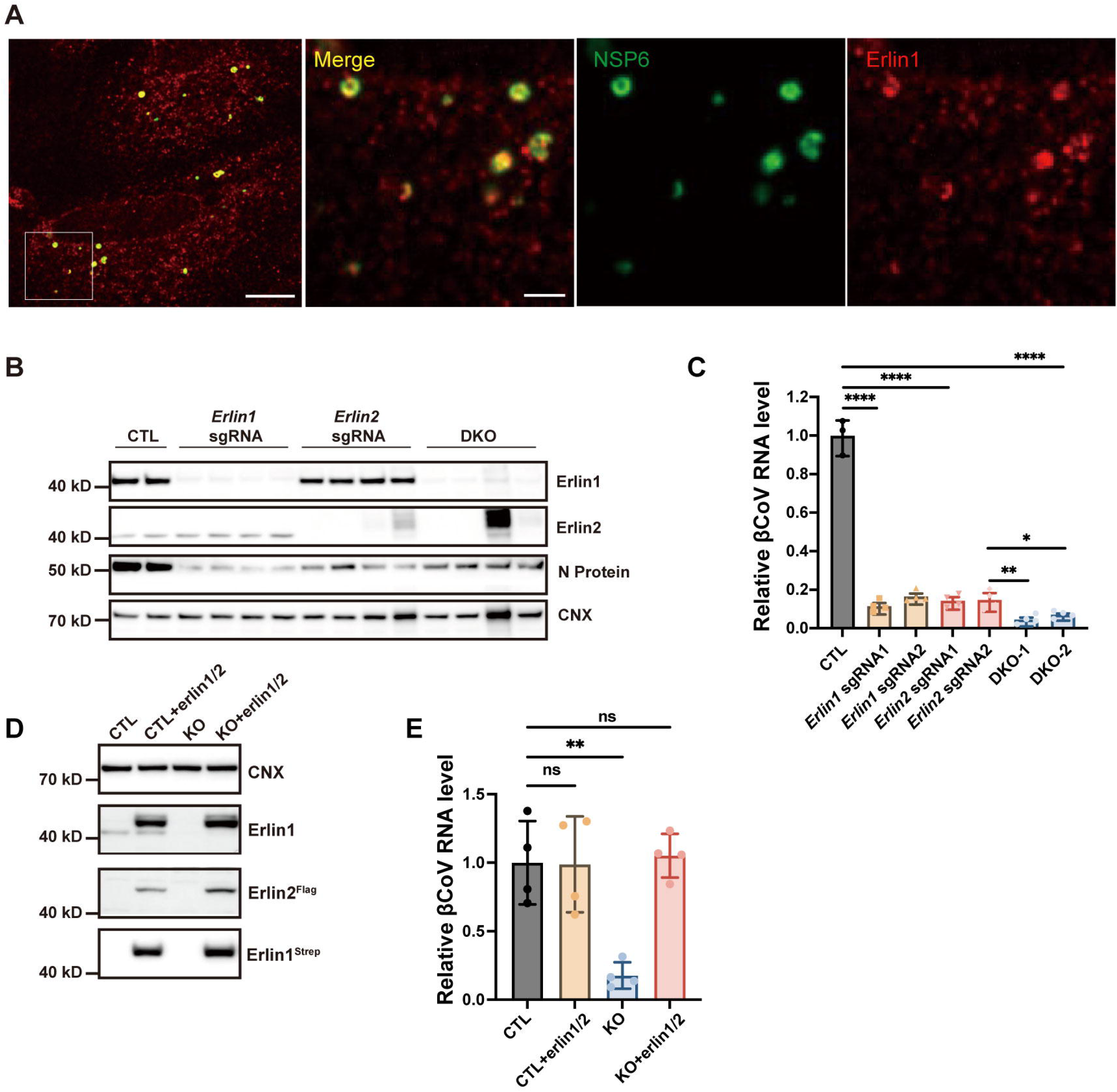
Functional requirement of erlin1/2 in β-coronavirus replication. (A) Co-localization of erlin1 with β-coronavirus (βCoV) replication-associated NSP6 protein. Whole-cell (left, scale bar: 10□μm) and zoomed-in (right, scale bar: 2□μm) views show the subcellular localizations of NSP6 and erlin1. mEGFP-tagged SARS-CoV-2 NSP6 (green) was overexpressed in A549 cells via AAV infection, and endogenous erlin1 (red) was detected by immunofluorescence staining using an erlin1 antibody. (B) Immunoblot (IB) analysis of βCoV-induced viral protein (N protein) expression in erlin1 and/or erlin2 knockout cells. βCoV-induced viral protein expression was diminished in A549-Cas9-mCEACAM1 cells upon CRISPR/Cas9-mediated single or double knockout of erlin1 and erlin2, using the indicated sgRNAs. (C) Erlin1 and/or erlin2 deficiency inhibits βCoV production in Hepa1-6 cells. CRISPR/Cas9-mediated erlin1 and/or erlin2 KO Hepa1-6 cells were infected with βCoV and harvested at 16 hr post-infection for RT-qPCR. Data represent the mean ± SEM from ≥3 independent experiments. ****p < 0.0001, **p < 0.01, *p < 0.05. (D) IB analysis of control (CTL) and erlin1 and erlin2 KO Hepa1-6 cells rescued with AAV-mediated erlin1 and erlin2 expression. (E) Rescue of βCoV inhibition in erlin1/2-deficient cells following erlin1/2 re-expression. Representative results from 4 independent experiments are shown. Data are presented as mean ± SEM. **p < 0.01, ns, not significant.

To examine the functional role of erlin1 and erlin2 in viral infection, we generated CRISPR/Cas9-mediated erlin1/2 knockout (KO) in human A549 cells bearing the MHV receptor mCEACAM1. Following infection with the β-coronavirus (MHV-A59), we found that knockout of erlin1 or erlin2 individually resulted in a marked decrease in the expression of the viral nucleocapsid (N) protein, a genome-associated structural component that reflects the level of viral RNA replication (Figure 6B). A similar inhibitory effect on viral RNA replication was observed following infection of erlin1/2 double-knockout murine Hepa1-6 cells, with a reduction exceeding 80%, whereas simultaneous knockout of both genes led to a reduction greater than 90% (Figure 6C). Importantly, reintroduction of erlin1/2 complex rescued the inhibitory phenotype of erlin KO and restored viral RNA replication levels (Figures 6D and 6E). Collectively, these findings firmly establish erlin1 and erlin2 as critical host factors required for β-coronavirus replication, potentially through their roles in organizing FMMs on the ER membrane.

## Discussion

Using cryo-electron microscopy, we determined that erlin1 and erlin2 co-assemble into a 26-mer cage-like structure, composed of alternating subunits and inserted into the ER membrane from the luminal side. This polymeric architecture is not strictly rigid and likely confers structural plasticity in response to membrane dynamics. In fact, on the raw cryo-EM images, various deformations could be observed on the erlin1/2 cages, suggesting that the membrane-attached erlin1/2 cages are intrinsically dynamic. The conserved SPFH1 domain potentially interacts with specific lipids and physically segregates the luminal leaflet of the ER membrane to form isolated microdomains. Notably, both the inner and outer surfaces of the erlin1/2 cages were capable of interacting with cargo proteins, suggesting that the erlin1/2 complex may function as a general scaffold for diverse ER-associated proteins.

### Organization of FMMs on the ER membrane by the erlin1/2 complex

SPFH family proteins are generally recognized as scaffold proteins that organize FMMs enriched in saturated lipids such as cholesterol and sphingomyelin^10,45^. However, the existence of such domains within the ER has remained controversial due to its relatively low cholesterol content. The identification of erlin1 and erlin2 — SPFH family members localized on the ER — provides compelling evidence supporting the presence of liquid-ordered (Lo) phase microdomains on the ER membrane^17^. Understanding how the erlin1/2 complex organizes these FMMs and regulates the proteins enriched within them represents a central focus of our study.

Through structural analysis, we found that erlin1 and erlin2 heteropolymerize to form cage-like complexes. The SPFH1 domain of the complex is embedded in the lumen-facing leaflet of the ER membrane, delineating a circular membrane region approximately 24 nm in diameter. This region exhibits resistance to solubilization by mild non-ionic detergents such as a mixture of GDN and CHS, suggesting an enrichment in saturated lipids within the erlin1/2 microdomain. We propose that this area represents the fundamental unit of ER-associated FMMs.

More importantly, the erlin1/2 cage may also regulate the lipid composition within the microdomain. The 26 SPFH1 domains tightly interact and completely seal off a circular area on the luminal leaflet of the ER membrane. Inside this microdomain, the diffusion and exchange of lipids in the luminal leaflet with the exterior membrane region is strictly prohibited, whereas the sparsely distributed 26 transmembrane helices in the cytoplasmic leaflet still permit the possible in-and-out movements of lipid molecules. The ER is the factory for biogenesis of most lipid species^46,47^. Compared to the plasma membrane, the ER is less asymmetrical in lipid composition between two leaflets^48^. Nevertheless, lines of evidence suggest that a transverse asymmetry in lipid composition is also essential for many biological processes occurring on the ER^46,49,50^. Our findings therefore suggest that erlin1/2 cages may also contribute to the maintenance of lipid asymmetry between the two leaflets within these microdomains. Importantly, we have identified a lipid binding pocket on the SPFH1 domain of erlin1/2, which could perfectly accommodate the head group of PI. Notably, PI can undergo phosphorylation and subsequent hydrolysis to generate second messengers such as inositol 1,4,5-trisphosphate (IP3) and diacylglycerol (DAG), which are critical mediators of signal transduction^51^. While PI is synthesized on the ER, its phosphorylation and dephosphorylation predominantly occur on the cytosolic leaflet of organelle membranes, serving as key determinants of membrane compartment identity^51,52^. The binding of erlin1/2 to PI may sequester it on the luminal side of the ER membrane, thereby preventing its phosphorylation and potentially disrupting the production or function of downstream signaling molecules.

Cryo-EM analysis further revealed that individual erlin1/2 cages can associate to form higher-order structures. In parallel, immunostaining of endogenous erlin1 showed its distribution in punctate structures of heterogeneous sizes on the ER membrane. These observations suggest that variable numbers of erlin1/2 complexes may assemble into higher-order clusters, ultimately contributing to the formation of FMMs of diverse sizes.

### Regulatory mechanisms of the erlin1/2 complex on its cargo proteins

In this study, we found that the erlin1/2 complex associates with a diverse array of cargo proteins. Is there a unified regulatory mechanism that governs all of these cargo proteins? We propose several hypotheses. For cargo proteins located within the cage, we suggest that the erlin1/2 complex may regulate them through two distinct mechanisms: (1) spatial sequestration— by enclosing cargo proteins through a sealed cage, the erlin1/2 complex would completely block the interaction between cargo proteins and their binding partners, either membrane or soluble proteins, from the luminal side (Figure 7A); and (2) lateral restriction — by partitioning the ER membrane into circular microdomains via the SPFH1 domains, erlin1/2 may physically separate membrane proteins, thereby limiting their proximity (Figure 7B). In both cases, the cytosol exposed regions of cargo proteins are still freely accessible from the cytosol; but the interactions between transmembrane domains of two proteins would also be limited. Given the presence of cracked cages in the reconstructed structures and various cage deformations on the raw cryo-EM particles (Figure 5A), we reason that these cage-associated regulations should be dynamic, reversible, and subject to further regulation through the assembly/disassembly cycle of the erlin1/2 cages.

**Figure 7.**
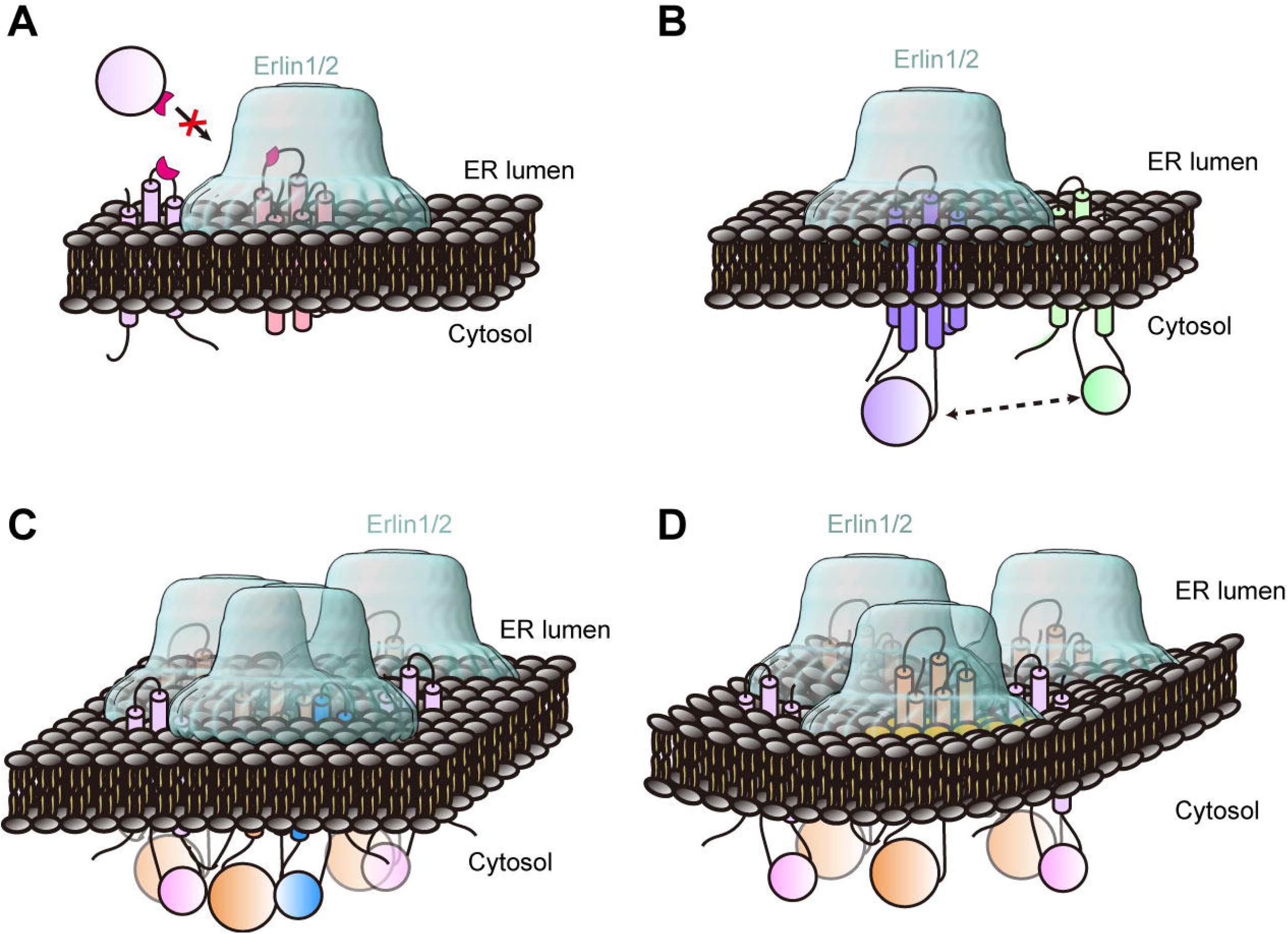
Proposed functional models of the erlin1/2 complex. (A) The erlin1/2 complex functions to spatially sequester membrane proteins on the ER, preventing their access from the luminal side. (B) The erlin1/2 complex functions to laterally restrict membrane proteins, limiting the interactions of their transmembrane or cytosol regions with their partners outside the cage. (C) The erlin1/2 complex increases the local concentration of cargo proteins through self-clustering, thereby enhancing cargo function via a concentration-dependent effect. Additionally, the erlin1/2 cluster may serve as a hub that brings cargo proteins and their associated partners into close proximity, increasing the likelihood of their interaction. (D) The erlin1/2 complex responds to certain stimuli to undergo conformational change, leading to the curvature change of the local membrane. Conversely, local membrane curvature changes induced by other cellular events may trigger conformational changes of the erlin1/2 cage, thereby regulating the activity of associated cargo proteins.

Furthermore, since erlin1/2 can organize FMMs on the ER membrane through cage clustering, we further speculate that it may serve as a functional platform for its cargo proteins that rely on their cytosolic domains for biological functions. Such platforms could enhance the local concentration of certain cargos, thereby amplifying their activity, and co-localize associated proteins within the same membrane region, increasing the likelihood of their interactions (Figure 7C). One such example is the ERAD process involving the ubiquitination of InsP3Rs. Previous studies have shown that RNAi-mediated knockdown^19,25,27^ or CRISPR-based knockout^26^ of erlin1/2 disrupt the ubiquitination and degradation of InsP3Rs.

Lastly, we propose that the erlin1/2 complex may regulate its cargo proteins by responding to specific stimuli and modulating local membrane curvature through cage expansion and contraction (Figure 7D). For instance, lipids synthesized on the cytoplasmic side of the ER membrane accumulate asymmetrically, causing the membrane to bulge toward the cytoplasm to accommodate this imbalance^50,53^, and may cause the erlin1/2 cage to deform, thereby altering the conformation and activity of proteins sequestered within the cage.

Collectively, these parallel mechanisms may act in concert to modulate the diffusion, concentration, activity, and interactions of erlin1/2 cargo proteins within spatially defined ER membrane territories. In summary, we have characterized the cage-like structure of the erlin1/2 complex in distinct conformational states and proposed a preliminary model of its regulatory mechanisms. These findings underscore a potential general role for erlin1/2 in the compartmentalization and functional modulation of the ER membrane environment.

## STAR⍰METHODS

### KEY RESOURCES TABLE

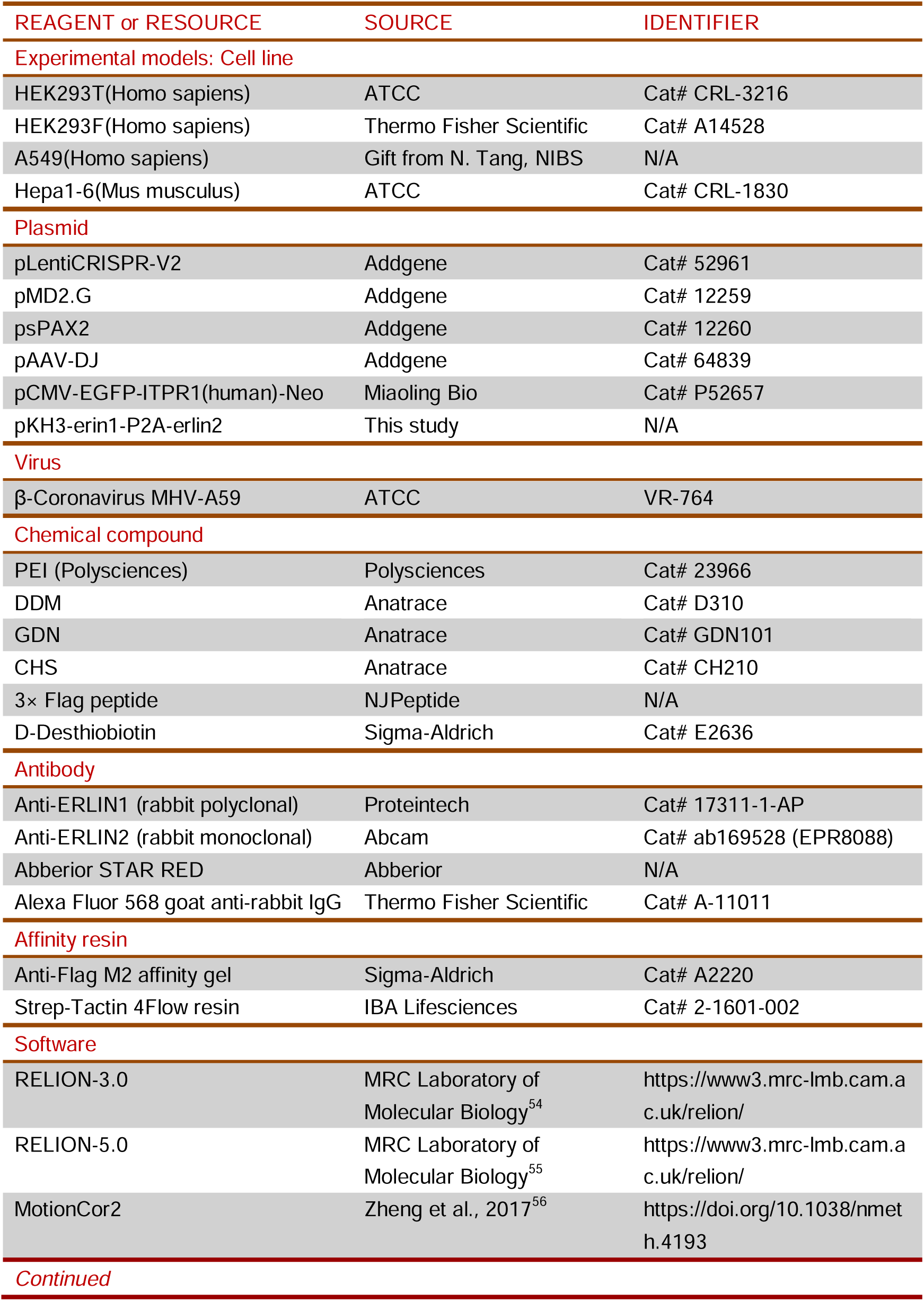
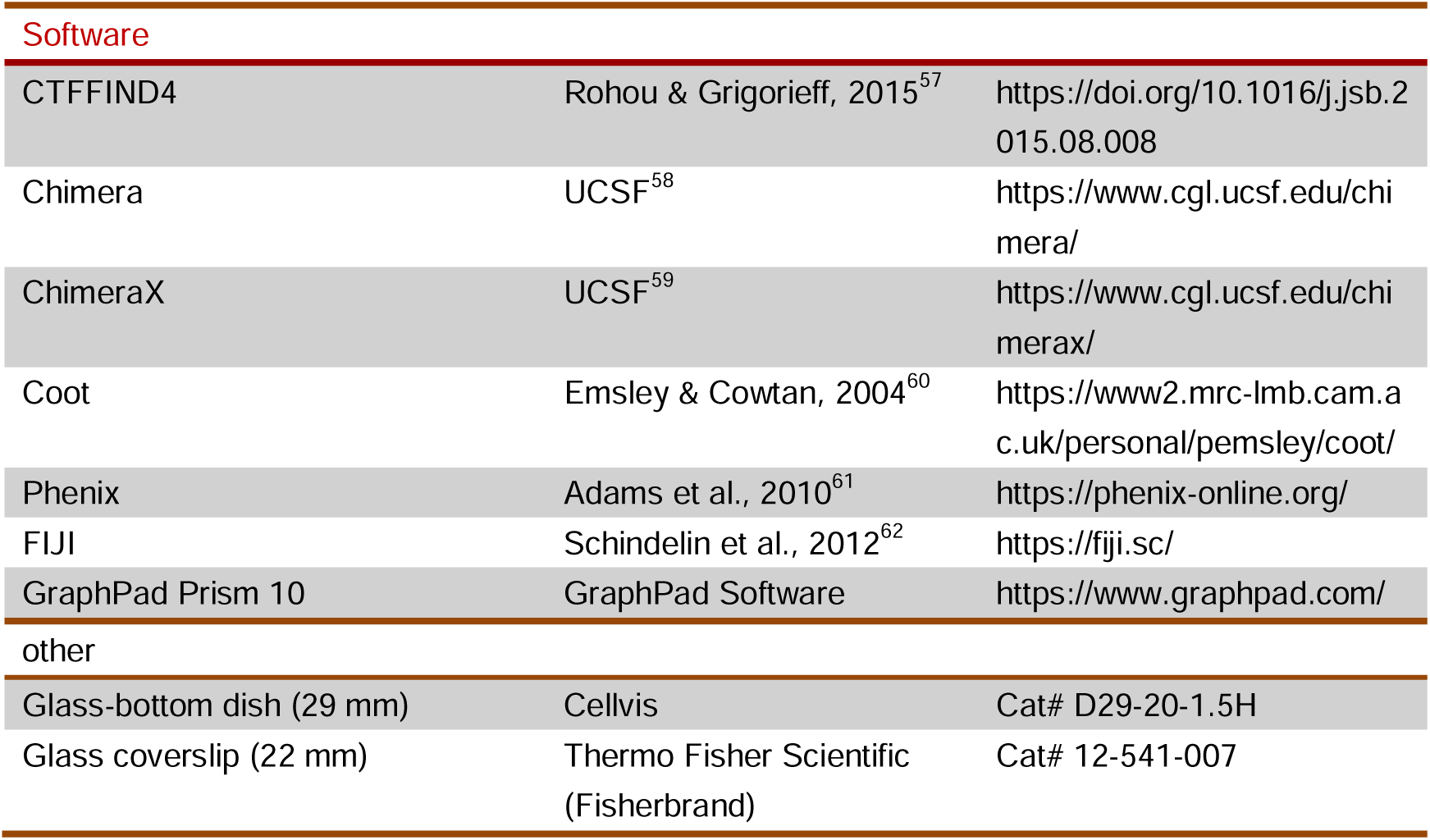

### EXPERIMENTAL MODEL AND STUDY PARTICIPANT DETAILS

#### Cell culture and treatments

HEK293T (ATCC, CRL-3216) and Hepa1-6 (ATCC, CRL-1830) cells were obtained from ATCC. A549 cells were generously provided by N. Tang (National Institute of Biological Sciences, Beijing). All cells are maintained in DMEM (Gibco) supplemented with 10% FBS and 1% penicillin/streptomycin at 37°C incubator with 5% CO2. HEK293F (Thermo Fisher Scientific, A14528) cells were cultured in SMM 293-TII (Sino Biological), in a 37□°C and 5% CO2 incubator shaking at 120 rpm.

### METHOD DETAILS

#### Cloning, expression and purification of the erlin1/2 complex

The genes of erlin1 and erlin2 were amplified from the cDNA library of HEK293F cells. The erlin1 gene was fused with a twin-Strep tag, and the erlin2 gene was fused with a Flag tag. The two genes were connected by P2A and cloned into pKH3 vector. Plasmid DNA and PEI (Polysciences) were transfected into HEK293F cells at a density of approximately 2×10□ cells/mL, using a mass ratio of 1:3 (plasmid:PEI). Transfection was performed with 1 mg of plasmid DNA per liter of culture.

Cells were harvested by centrifugation and washed once with PBS. The cell pellet was resuspended in lysis buffer (40 mM HEPES-KOH, pH 8.0, 150 mM NaCl, 10% glycerol) supplemented with 1× cocktail protease inhibitor (Mei5bio) and lysed by sonication. The lysate was centrifuged at 1,300 × *g* for 15 min to remove cell debris, and the supernatant was centrifuged at 140,000 × *g* for 1 hr in a 70Ti rotor (Beckman Coulter) to collect the membrane fraction. The pelleted membranes were solubilized in lysis buffer containing 0.75% (w/v) n-dodecyl-β-d-maltoside (DDM, Anatrace) and 0.25% (w/v) GDN (Anatrace) or 1% (w/v) GDN (Anatrace) and 0.1% (w/v) cholesterol hemisuccinate (CHS, Anatrace) at 4°C for 4 hr. The supernatant was collected by centrifugation at 170,000 × *g* for 30 min in a 70Ti rotor (Beckman Coulter). The supernatant was incubated with anti-Flag M2 affinity resin (Sigma Aldrich) or Strep-Tactin 4Flow (iba) at 4°C for 2 hr with rotation, followed by washing with 100 c.v. of wash buffer (lysis buffer supplemented with 1× cocktail protease inhibitor and 0.042% (w/v) GDN) and elution with wash buffer containing 0.5 mg/mL 3× Flag peptide (NJPeptide) or 20 mM d-Desthiobiotin (Sigma-Aldrich). The eluate was concentrated with a 100-kDa cutoff spin concentrator (Merck Millipore) and loaded onto a 10%–50% glycerol density gradient and centrifuged at 100,000 × *g* for 14 hr in a SW41 Ti rotor (Beckman Coulter). All fractions of the gradient were collected and analyzed by SDS-PAGE. The peak fraction was concentrated for cryo-EM sample preparation.

The purification of the erlin mutant was performed using a TLS-55 rotor (Beckman Coulter) for density gradient centrifugation, with all other conditions remaining the same.

#### Negative staining electron microscopy examination

4 uL of sample at a concentration of approximately 0.2 mg/mL was loaded onto a copper grid and stained with 2% uranyl acetate. The grid was examined using a JEOL JEM-F200 at 200 kV.

#### Cryo-EM sample preparation and data collection

4 μL of sample with a concentration of approximately 10 mg/mL was loaded onto a glow-discharged gold grid (Quantifoil, R2/1, 300 mesh), waited for 4 s, and blotted for 1□s with –1 blot force using an FEI Vitrobot at 6□°C and 100% humidity.

Data (DDM+GDN) were collected using a FEI Titan Krios (with Gatan K2 summit camera). Movies were collected using SerialEM^63^ at 130,000× magnification (pixel size of 1.052□Å) with the defocus range varying from −1.2 μm to −1.8 μm. Each movie contained 32 frames with a dose rate of 8 e^−^/Å^2^/s for a total exposure time of 8□s.

Data (GDN+CHS) were collected using a Thermo Fisher Scientific Krios G4 (with Falcon 4 camera). Movies were collected using EPU^64^ at 130,000× magnification (pixel size of 0.95□Å) with the defocus range varying from −1.2 μm to −1.8 μm. Each movie contained 32 frames with a dose rate of 9.84 e^−^/Å^2^/s for a total exposure time of 5.5□s.

#### Cryo-EM data processing

The motion correction and electron-dose weighting of the movies were performed with MotionCor2^56^. The CTF parameters were estimated with the program of CTFfind4^57^. Particle picking, extraction, classification and refinement were done with RELION-3.0^54^ and RELION-5.0^55^.

The particles (DDM+GDN) selected by the Laplace-Gaussian algorithm were subjected to multiple rounds of 2D classification and 3D classification. The good classes were selected for 3D refinement with C13 symmetry. The particle chirality is separated by 3D classification, and the chiral density map is flipped. After merging the particles, two rounds of 3D refinement with C13 symmetry, one round of 3D refinement with C26 symmetry, and then another round of 3D refinement with C13 symmetry were performed to improve the overall resolution. For the calculation of the C-terminus with poor resolution, the center of the particle is moved to the C-terminus, 3D classification (skip alignment) is performed, and several good classes are selected for mask-based 3D refinement.

The particles (GDN+ CHS) selected by the Laplace-Gaussian algorithm were subjected to multiple rounds of 2D classification. 42,938 particles from the good classes were selected for Topaz training and selection. The selected particles were subjected to multiple rounds of 2D and 3D classification. The selected particles were subjected to 3D refinement with C13 symmetry. In order to further improve the overall resolution, the refinement was performed with a mask-based 3D classification (skip alignment) and multiple states were separated. In order to further improve the C-terminal resolution, the particle center was moved and another mask-based 3D classification (skip alignment) was performed.

#### Model building

The monomer structures of erlin1 and erlin2 were generated by Alphafold protein structure database (https://alphafold.ebi.ac.uk). Models were fitted into the maps by rigid-body fitting in UCSF Chimera^58^ and ChimeraX^59^, followed by manual adjustment in Coot^60^. The construction of the C-terminus model uses the local map generated by particle center shifting. Model refinement was done using real-space refinement in Phenix^61^. Models were evaluated in MolProbity^65^. The figures preparation and structure analysis were performed with Chimera and ChimeraX.

#### Co-immunoprecipitation Assay

The genes encoding erlin1/2 and EGFP tagged InsP3R1 (P52657, Miaolingbio) were co-expressed in HEK293F cells. Membrane proteins were extracted from an equal number of cells using either 0.75% (w/v) DDM and 0.25% (w/v) GDN or 1% (w/v) GDN and 0.1% (w/v) CHS. The supernatant was collected by centrifugation at 186,000 × *g* for 10 min in a TLA-55 rotor (Beckman Coulter). An equal volume of supernatant was incubated with either anti-Flag M2 affinity resin or Streptactin Beads 4FF, and purified as described previously. Bound proteins were denatured directly in 4× SDS loading buffer at 37°C for 30 min. All samples were analyzed by SDS-PAGE followed by Western blotting.

#### Immunofluorescence staining and STED imaging analysis

U2-OS cells were cultured on 29 mm glass-bottom dishes (Cellvis, D29-20-1.5H) for treatment and immunofluorescence staining. Cells were washed three times with pre-warmed PBS (37°C) and fixed with 4% paraformaldehyde (PFA) for 15 min at room temperature, followed by three additional PBS washes. The cell membrane was then permeabilized with 0.1% Triton X-100. Non-specific binding sites were blocked by incubating cells with 4% normal BSA in PBS for 1 hr at room temperature. Primary antibody incubation was performed overnight at 4°C using anti-ERLIN1 polyclonal antibody (Proteintech, 17311-1-AP) or anti-ERLIN2 antibody (Abcam, EPR8088) diluted in blocking solution. After three washes with PBS, cells were incubated with Abberior STAR RED-conjugated goat anti-rabbit IgG (H+L) cross-adsorbed secondary antibody (Abberior) for 40 min at room temperature, followed by three additional PBS washes.

Fluorescence images were acquired using a Leica TCS SP8 STED 3X microscope. For statistical analysis, images were uniformly acquired at a size of 23.25 μm × 23.25 μm with a pixel resolution of 22.73 nm × 22.73 nm. The collected images were processed using FIJI software^62^, where the diameter and number of puncta structures were quantified. The data were classified into 50 nm bins, and statistical graphs were generated using GraphPad Prism 10.

#### AAV Packaging, Transduction, and Immunofluorescence Microscopy

AAV packaging and purification were performed as previously described^66^. Briefly, HEK293T cells were transfected with AAV shuttle plasmids, pAAV-DJ, and helper plasmids using polyethylenimine (PEI). At 60 hours post-transfection, cells were harvested, and viral particles were purified and quantified by Coomassie Brilliant Blue R250 staining and quantitative PCR (qPCR). For gene delivery, cultured cells in 6-well plates were transduced with AAV-DJ at a dose of 1 × 10¹□ genome copies per well. Rescue experiments were conducted 7 days post-infection, and immunofluorescence assays were performed 36 hours after AAV-DJ infection.

For immunofluorescence analysis, cells were cultured on 22-mm diameter glass coverslips (Fisherbrand_™_, Thermo Fisher Scientific, Cat# 12-541-007) prior to treatment and staining.Cells were fixed in cold methanol for 5 minutes at room temperature, followed by three washes with phosphate-buffered saline (PBS). Non-specific binding was blocked using 2.5% normal goat serum in PBS for 1 hour at room temperature. Primary antibody incubation was performed overnight at 4°C using rabbit anti-ERLIN1 antibody (Proteintech, Cat# 17311-1-AP; 1:200 dilution) diluted in blocking buffer. After washing three times with PBS, cells were incubated for 1 hour at room temperature with Alexa Fluor_™_ 568-conjugated goat anti-rabbit IgG (H+L), cross-adsorbed secondary antibody (Thermo Fisher Scientific, Cat# A-11011; 1:1000 dilution), followed by three additional PBS washes.

Fluorescence images were acquired using a ZEISS spinning disk confocal microscope and processed using ImageJ software (National Institutes of Health).

#### Construction of the erlin1/2 knockout cell lines and **β**CoV infection experiments

Single-guide RNAs (sgRNAs) targeting erlin1, erlin2, and Lacz (control) were designed using Benchling (https://benchling.com/) to ensure high editing efficiency and minimal off-target effects (Supplementary Table 1). These sgRNAs were cloned into the pLentiCRISPR-V2 vector for lentiviral delivery. Lentiviruses were produced in HEK293T cells by co-transfection with psPAX2 and pMD2.G packaging plasmids using PEI. Supernatants containing lentivirus were collected 48 hr post-transfection and used to transduce wild-type Hepa1-6 and A549-Cas9-mCEACAM1 cells. Two days after infection, cells were selected with appropriate antibiotics to generate erlin1 and erlin2 knockout cell lines.

For β-coronavirus (MHV-A59) infection, viral stocks were prepared in 17Cl-1 cells and quantified by plaque assay. A549-Cas9-mCEACAM1 cells were infected at MOI = 0.005, and Hepa1-6 cells at MOI = 0.2. After 1 hr of virus adsorption, the medium was replaced with DMEM supplemented with 10% FBS and 1% penicillin/streptomycin. Cells were collected at 8 (A549-Cas9-mCEACAM1) or 16 (Hepa1-6) hr post-infection for RNA extraction and RT-qPCR, and at 24 hr for immunoblotting.

Total RNA was isolated from cell lysates using Trizol reagent according to the manufacturer’s protocol. cDNA synthesis was performed using the SuperScript III Reverse Transcriptase kit (TransGen Biotech) with oligo(dT) primers, following the manufacturer’s instructions. Quantitative real-time PCR (qPCR) was carried out using SYBR Green Master Mix on an Archimed X4 fluorescence quantification system (Rocgene). All primer sequences used in this study are provided in Supplementary Table 2.

## Acknowledgements

We thank the Core Facilities at the School of Life Sciences, Peking University (PKU) for help with negative staining EM; the PKU Cryo-EM Platform and the Electron Microscopy Laboratory of PKU for help with data collection; the High-performance Computing Platform of Peking University for help with computation; and the National Center for Protein Sciences at Peking University for assistance with imaging. This work was supported by National Natural Science Foundation of China (92354306 to N.G.).

## Author contributions

L.Y. and Z.X. prepared the samples of the erlin1/2 complex and collected the cryo-EM data. L.Y. and Z.X. performed EM analysis (with the help of N.L., and C.M.). N.G. and L.Y. performed cryo-EM model building. Z.X., L.Y. and X.W. performed co-IP experiments and endogenous erlin1/2 Immunofluorescence imaging. Y.Y. established the CRISPR KO cell lines and performed the βCoV infection experiments (with the help of Y.W.). N.G., L.Y., Z.X. and Y.Y. wrote the manuscript. N.G. and X.C. supervised the study.

## Data and materials availability

The cryo-EM map and the structure coordinates have been deposited in the Electron Microscopy Data Bank and the Protein Data Bank under accession codes EMD-XXXX and XXXX.

## Supplementary Materials

Figures. S1 to S9

Table S1 to S3

## References

1. Lingwood, D., and Simons, K. (2010). Lipid rafts as a membrane-organizing principle. Science 327, 46–50. 10.1126/science.1174621.

2. Sezgin, E., Levental, I., Mayor, S., and Eggeling, C. (2017). The mystery of membrane organization: composition, regulation and roles of lipid rafts. Nat Rev Mol Cell Biol 18, 361–374. 10.1038/nrm.2017.16.

3. Munro, S. (2003). Lipid rafts: elusive or illusive? Cell 115, 377–388. 10.1016/s0092-8674(03)00882-1.

4. Simons, K., and Ikonen, E. (1997). Functional rafts in cell membranes. Nature 387, 569–572. 10.1038/42408.

5. Subczynski, W.K., and Kusumi, A. (2003). Dynamics of raft molecules in the cell and artificial membranes: approaches by pulse EPR spin labeling and single molecule optical microscopy. Biochim Biophys Acta 1610, 231–243. 10.1016/s0005-2736(03)00021-x.

6. Kusumi, A., and Suzuki, K. (2005). Toward understanding the dynamics of membrane-raft-based molecular interactions. Biochim Biophys Acta 1746, 234–251. 10.1016/j.bbamcr.2005.10.001.

7. Brown, D.A. (2007). Analysis of raft affinity of membrane proteins by detergent-insolubility. Methods Mol Biol 398, 9–20. 10.1007/978-1-59745-513-8_2.

8. Field, K.A., Holowka, D., and Baird, B. (1995). Fc epsilon RI-mediated recruitment of p53/56lyn to detergent-resistant membrane domains accompanies cellular signaling. Proc Natl Acad Sci U S A 92, 9201–9205. 10.1073/pnas.92.20.9201.

9. Yokoyama, H., and Matsui, I. (2020). The lipid raft markers stomatin, prohibitin, flotillin, and HflK/C (SPFH)-domain proteins form an operon with NfeD proteins and function with apolar polyisoprenoid lipids. Crit Rev Microbiol 46, 38–48. 10.1080/1040841X.2020.1716682.

10. Browman, D.T., Hoegg, M.B., and Robbins, S.M. (2007). The SPFH domain-containing proteins: more than lipid raft markers. Trends Cell Biol 17, 394–402. 10.1016/j.tcb.2007.06.005.

11. Browman, D.T., Resek, M.E., Zajchowski, L.D., and Robbins, S.M. (2006). Erlin-1 and erlin-2 are novel members of the prohibitin family of proteins that define lipid-raft-like domains of the ER. J Cell Sci 119, 3149–3160. 10.1242/jcs.03060.

12. Stewart, G.W. (1997). Stomatin. Int J Biochem Cell Biol 29, 271–274. 10.1016/s1357-2725(96)00072-6.

13. Lapatsina, L., Brand, J., Poole, K., Daumke, O., and Lewin, G.R. (2012). Stomatin-domain proteins. Eur J Cell Biol 91, 240–245. 10.1016/j.ejcb.2011.01.018.

14. Steglich, G., Neupert, W., and Langer, T. (1999). Prohibitins regulate membrane protein degradation by the m-AAA protease in mitochondria. Mol Cell Biol 19, 3435–3442. 10.1128/MCB.19.5.3435.

15. Bickel, P.E., Scherer, P.E., Schnitzer, J.E., Oh, P., Lisanti, M.P., and Lodish, H.F. (1997). Flotillin and epidermal surface antigen define a new family of caveolae-associated integral membrane proteins. J Biol Chem 272, 13793–13802. 10.1074/jbc.272.21.13793.

16. Schwarz, K., Simons, M., Reiser, J., Saleem, M.A., Faul, C., Kriz, W., Shaw, A.S., Holzman, L.B., and Mundel, P. (2001). Podocin, a raft-associated component of the glomerular slit diaphragm, interacts with CD2AP and nephrin. J Clin Invest 108, 1621–1629. 10.1172/JCI12849.

17. Manganelli, V., Longo, A., Mattei, V., Recalchi, S., Riitano, G., Caissutti, D., Capozzi, A., Sorice, M., Misasi, R., and Garofalo, T. (2021). Role of ERLINs in the Control of Cell Fate through Lipid Rafts. Cells 10. 10.3390/cells10092408.

18. Pearce, M.M.P., Wormer, D.B., Wilkens, S., and Wojcikiewicz, R.J.H. (2009). An Endoplasmic Reticulum (ER) Membrane Complex Composed of SPFH1 and SPFH2 Mediates the ER-associated Degradation of Inositol 1,4,5-Trisphosphate Receptors. Journal of Biological Chemistry 284, 10433–10445. 10.1074/jbc.M809801200.

19. Pearce, M.M., Wang, Y., Kelley, G.G., and Wojcikiewicz, R.J. (2007). SPFH2 mediates the endoplasmic reticulum-associated degradation of inositol 1,4,5-trisphosphate receptors and other substrates in mammalian cells. J Biol Chem 282, 20104–20115. 10.1074/jbc.M701862200.

20. Lee, J.N., Song, B., DeBose-Boyd, R.A., and Ye, J. (2006). Sterol-regulated degradation of Insig-1 mediated by the membrane-bound ubiquitin ligase gp78. J Biol Chem 281, 39308–39315. 10.1074/jbc.M608999200.

21. Vembar, S.S., and Brodsky, J.L. (2008). One step at a time: endoplasmic reticulum-associated degradation. Nat Rev Mol Cell Biol 9, 944–957. 10.1038/nrm2546.

22. Lemberg, M.K., and Strisovsky, K. (2021). Maintenance of organellar protein homeostasis by ER-associated degradation and related mechanisms. Mol Cell 81, 2507–2519. 10.1016/j.molcel.2021.05.004.

23. Wright, F.A., Lu, J.P., Sliter, D.A., Dupre, N., Rouleau, G.A., and Wojcikiewicz, R.J. (2015). A Point Mutation in the Ubiquitin Ligase RNF170 That Causes Autosomal Dominant Sensory Ataxia Destabilizes the Protein and Impairs Inositol 1,4,5-Trisphosphate Receptor-mediated Ca2+ Signaling. J Biol Chem 290, 13948–13957. 10.1074/jbc.M115.655043.

24. Wojcikiewicz, R.J., Pearce, M.M., Sliter, D.A., and Wang, Y. (2009). When worlds collide: IP(3) receptors and the ERAD pathway. Cell Calcium 46, 147–153. 10.1016/j.ceca.2009.05.002.

25. Wang, Y., Pearce, M.M., Sliter, D.A., Olzmann, J.A., Christianson, J.C., Kopito, R.R., Boeckmann, S., Gagen, C., Leichner, G.S., Roitelman, J., and Wojcikiewicz, R.J. (2009). SPFH1 and SPFH2 mediate the ubiquitination and degradation of inositol 1,4,5-trisphosphate receptors in muscarinic receptor-expressing HeLa cells. Biochim Biophys Acta 1793, 1710–1718. 10.1016/j.bbamcr.2009.09.004.

26. Wright, F.A., Bonzerato, C.G., Sliter, D.A., and Wojcikiewicz, R.J.H. (2018). The erlin2 T65I mutation inhibits erlin1/2 complex-mediated inositol 1,4,5-trisphosphate receptor ubiquitination and phosphatidylinositol 3-phosphate binding. J Biol Chem 293, 15706–15714. 10.1074/jbc.RA118.004547.

27. Lu, J.P., Wang, Y., Sliter, D.A., Pearce, M.M., and Wojcikiewicz, R.J. (2011). RNF170 protein, an endoplasmic reticulum membrane ubiquitin ligase, mediates inositol 1,4,5-trisphosphate receptor ubiquitination and degradation. J Biol Chem 286, 24426–24433. 10.1074/jbc.M111.251983.

28. Jo, Y., Sguigna, P.V., and DeBose-Boyd, R.A. (2011). Membrane-associated ubiquitin ligase complex containing gp78 mediates sterol-accelerated degradation of 3-hydroxy-3-methylglutaryl-coenzyme A reductase. J Biol Chem 286, 15022–15031. 10.1074/jbc.M110.211326.

29. Huber, M.D., Vesely, P.W., Datta, K., and Gerace, L. (2013). Erlins restrict SREBP activation in the ER and regulate cellular cholesterol homeostasis. J Cell Biol 203, 427–436. 10.1083/jcb.201305076.

30. Wakil, S.M., Bohlega, S., Hagos, S., Baz, B., Al Dossari, H., Ramzan, K., and Al-Hassnan, Z.N. (2013). A novel splice site mutation in ERLIN2 causes hereditary spastic paraplegia in a Saudi family. Eur J Med Genet 56, 43–45. 10.1016/j.ejmg.2012.10.003.

31. Tian, W.T., Shen, J.Y., Liu, X.L., Wang, T., Luan, X.H., Zhou, H.Y., Chen, S.D., Huang, X.J., and Cao, L. (2016). Novel Mutations in Endoplasmic Reticulum Lipid Raft-associated Protein 2 Gene Cause Pure Hereditary Spastic Paraplegia Type 18. Chin Med J (Engl) 129, 2759–2761. 10.4103/0366-6999.193444.

32. Tunca, C., Akcimen, F., Coskun, C., Gundogdu-Eken, A., Kocoglu, C., Cevik, B., Bekircan-Kurt, C.E., Tan, E., and Basak, A.N. (2018). ERLIN1 mutations cause teenage-onset slowly progressive ALS in a large Turkish pedigree. Eur J Hum Genet 26, 745–748. 10.1038/s41431-018-0107-5.

33. Rydning, S.L., Dudesek, A., Rimmele, F., Funke, C., Kruger, S., Biskup, S., Vigeland, M.D., Hjorthaug, H.S., Sejersted, Y., Tallaksen, C., et al. (2018). A novel heterozygous variant in ERLIN2 causes autosomal dominant pure hereditary spastic paraplegia. Eur J Neurol 25, 943–e971. 10.1111/ene.13625.

34. Hoegg, M.B., Browman, D.T., Resek, M.E., and Robbins, S.M. (2009). Distinct regions within the erlins are required for oligomerization and association with high molecular weight complexes. J Biol Chem 284, 7766–7776. 10.1074/jbc.M809127200.

35. Solis, G.P., Hoegg, M., Munderloh, C., Schrock, Y., Malaga-Trillo, E., Rivera-Milla, E., and Stuermer, C.A. (2007). Reggie/flotillin proteins are organized into stable tetramers in membrane microdomains. Biochem J 403, 313–322. 10.1042/BJ20061686.

36. Kasashima, K., Ohta, E., Kagawa, Y., and Endo, H. (2006). Mitochondrial functions and estrogen receptor-dependent nuclear translocation of pleiotropic human prohibitin 2. J Biol Chem 281, 36401–36410. 10.1074/jbc.M605260200.

37. Ma, C.Y., Wang, C.K., Luo, D.Y., Yan, L., Yang, W.X., Li, N.N., and Gao, N. (2022). Structural insights into the membrane microdomain organization by SPFH family proteins. Cell Res 32, 176–189. 10.1038/s41422-021-00598-3.

38. Qiao, Z., Yokoyama, T., Yan, X.F., Beh, I.T., Shi, J., Basak, S., Akiyama, Y., and Gao, Y.G. (2022). Cryo-EM structure of the entire FtsH-HflKC AAA protease complex. Cell Rep 39, 110890. 10.1016/j.celrep.2022.110890.

39. Fu, Z., and MacKinnon, R. (2024). Structure of the flotillin complex in a native membrane environment. Proc Natl Acad Sci U S A 121, e2409334121. 10.1073/pnas.2409334121.

40. Abramson, J., Adler, J., Dunger, J., Evans, R., Green, T., Pritzel, A., Ronneberger, O., Willmore, L., Ballard, A.J., Bambrick, J., et al. (2024). Accurate structure prediction of biomolecular interactions with AlphaFold 3. Nature 630, 493–500. 10.1038/s41586-024-07487-w.

41. Inoue, T., and Tsai, B. (2013). How viruses use the endoplasmic reticulum for entry, replication, and assembly. Cold Spring Harb Perspect Biol 5, a013250. 10.1101/cshperspect.a013250.

42. Ci, Y., and Shi, L. (2021). Compartmentalized replication organelle of flavivirus at the ER and the factors involved. Cell Mol Life Sci 78, 4939–4954. 10.1007/s00018-021-03834-6.

43. Wolff, G., Melia, C.E., Snijder, E.J., and Barcena, M. (2020). Double-Membrane Vesicles as Platforms for Viral Replication. Trends Microbiol 28, 1022–1033. 10.1016/j.tim.2020.05.009.

44. Ricciardi, S., Guarino, A.M., Giaquinto, L., Polishchuk, E.V., Santoro, M., Di Tullio, G., Wilson, C., Panariello, F., Soares, V.C., Dias, S.S.G., et al. (2022). The role of NSP6 in the biogenesis of the SARS-CoV-2 replication organelle. Nature 606, 761–768. 10.1038/s41586-022-04835-6.

45. Langhorst, M.F., Reuter, A., and Stuermer, C.A. (2005). Scaffolding microdomains and beyond: the function of reggie/flotillin proteins. Cell Mol Life Sci 62, 2228–2240. 10.1007/s00018-005-5166-4.

46. Holthuis, J.C., and Menon, A.K. (2014). Lipid landscapes and pipelines in membrane homeostasis. Nature 510, 48–57. 10.1038/nature13474.

47. Jacquemyn, J., Cascalho, A., and Goodchild, R.E. (2017). The ins and outs of endoplasmic reticulum-controlled lipid biosynthesis. EMBO Rep 18, 1905–1921. 10.15252/embr.201643426.

48. van Meer, G., Voelker, D.R., and Feigenson, G.W. (2008). Membrane lipids: where they are and how they behave. Nat Rev Mol Cell Biol 9, 112–124. 10.1038/nrm2330.

49. Kobayashi, T., and Menon, A.K. (2018). Transbilayer lipid asymmetry. Curr Biol 28, R386–R391. 10.1016/j.cub.2018.01.007.

50. Chorlay, A., Monticelli, L., Verissimo Ferreira, J., Ben M’barek, K., Ajjaji, D., Wang, S., Johnson, E., Beck, R., Omrane, M., Beller, M., et al. (2019). Membrane Asymmetry Imposes Directionality on Lipid Droplet Emergence from the ER. Dev Cell 50, 25–42 e27. 10.1016/j.devcel.2019.05.003.

51. Balla, T. (2013). Phosphoinositides: tiny lipids with giant impact on cell regulation. Physiol Rev 93, 1019–1137. 10.1152/physrev.00028.2012.

52. Posor, Y., Jang, W., and Haucke, V. (2022). Phosphoinositides as membrane organizers. Nature Reviews Molecular Cell Biology 23, 797–816. 10.1038/s41580-022-00490-x.

53. Peeters, B.W.A., Piet, A.C.A., and Fornerod, M. (2022). Generating Membrane Curvature at the Nuclear Pore: A Lipid Point of View. Cells 11. 10.3390/cells11030469.

54. Zivanov, J., Nakane, T., Forsberg, B.O., Kimanius, D., Hagen, W.J., Lindahl, E., and Scheres, S.H. (2018). New tools for automated high-resolution cryo-EM structure determination in RELION-3. Elife 7. 10.7554/eLife.42166.

55. Burt, A., Toader, B., Warshamanage, R., von Kugelgen, A., Pyle, E., Zivanov, J., Kimanius, D., Bharat, T.A.M., and Scheres, S.H.W. (2024). An image processing pipeline for electron cryo-tomography in RELION-5. FEBS Open Bio 14, 1788–1804. 10.1002/2211-5463.13873.

56. Zheng, S.Q., Palovcak, E., Armache, J.P., Verba, K.A., Cheng, Y., and Agard, D.A. (2017). MotionCor2: anisotropic correction of beam-induced motion for improved cryo-electron microscopy. Nat Methods 14, 331–332. 10.1038/nmeth.4193.

57. Rohou, A., and Grigorieff, N. (2015). CTFFIND4: Fast and accurate defocus estimation from electron micrographs. J Struct Biol 192, 216–221. 10.1016/j.jsb.2015.08.008.

58. Pettersen, E.F., Goddard, T.D., Huang, C.C., Couch, G.S., Greenblatt, D.M., Meng, E.C., and Ferrin, T.E. (2004). UCSF Chimera--a visualization system for exploratory research and analysis. J Comput Chem 25, 1605–1612. 10.1002/jcc.20084.

59. Pettersen, E.F., Goddard, T.D., Huang, C.C., Meng, E.C., Couch, G.S., Croll, T.I., Morris, J.H., and Ferrin, T.E. (2021). UCSF ChimeraX: Structure visualization for researchers, educators, and developers. Protein Sci 30, 70–82. 10.1002/pro.3943.

60. Emsley, P., and Cowtan, K. (2004). Coot: model-building tools for molecular graphics. Acta Crystallogr D Biol Crystallogr 60, 2126–2132. 10.1107/S0907444904019158.

61. Adams, P.D., Afonine, P.V., Bunkoczi, G., Chen, V.B., Davis, I.W., Echols, N., Headd, J.J., Hung, L.W., Kapral, G.J., Grosse-Kunstleve, R.W., et al. (2010). PHENIX: a comprehensive Python-based system for macromolecular structure solution. Acta Crystallogr D Biol Crystallogr 66, 213–221. 10.1107/S0907444909052925.

62. Schindelin, J., Arganda-Carreras, I., Frise, E., Kaynig, V., Longair, M., Pietzsch, T., Preibisch, S., Rueden, C., Saalfeld, S., Schmid, B., et al. (2012). Fiji: an open-source platform for biological-image analysis. Nat Methods 9, 676–682. 10.1038/nmeth.2019.

63. Mastronarde, D.N. (2005). Automated electron microscope tomography using robust prediction of specimen movements. J Struct Biol 152, 36–51. 10.1016/j.jsb.2005.07.007.

64. Thompson, R.F., Iadanza, M.G., Hesketh, E.L., Rawson, S., and Ranson, N.A. (2019). Collection, pre-processing and on-the-fly analysis of data for high-resolution, single-particle cryo-electron microscopy. Nat Protoc 14, 100–118. 10.1038/s41596-018-0084-8.

65. Williams, C.J., Headd, J.J., Moriarty, N.W., Prisant, M.G., Videau, L.L., Deis, L.N., Verma, V., Keedy, D.A., Hintze, B.J., Chen, V.B., et al. (2018). MolProbity: More and better reference data for improved all-atom structure validation. Protein Sci 27, 293–315. 10.1002/pro.3330.

66. Wang, X., Xu, B.L., and Chen, X.W. (2021). Acute gene inactivation in the adult mouse liver using the CRISPR-Cas9 technology. STAR Protoc 2, 100611. 10.1016/j.xpro.2021.100611.

