## Supplementary material for "Erlin1/2 Complex is a Dynamic Scaffold for Membrane Protein Sequestration and Microdomain Assembly on the Endoplasmic Reticulum": Supplymentary

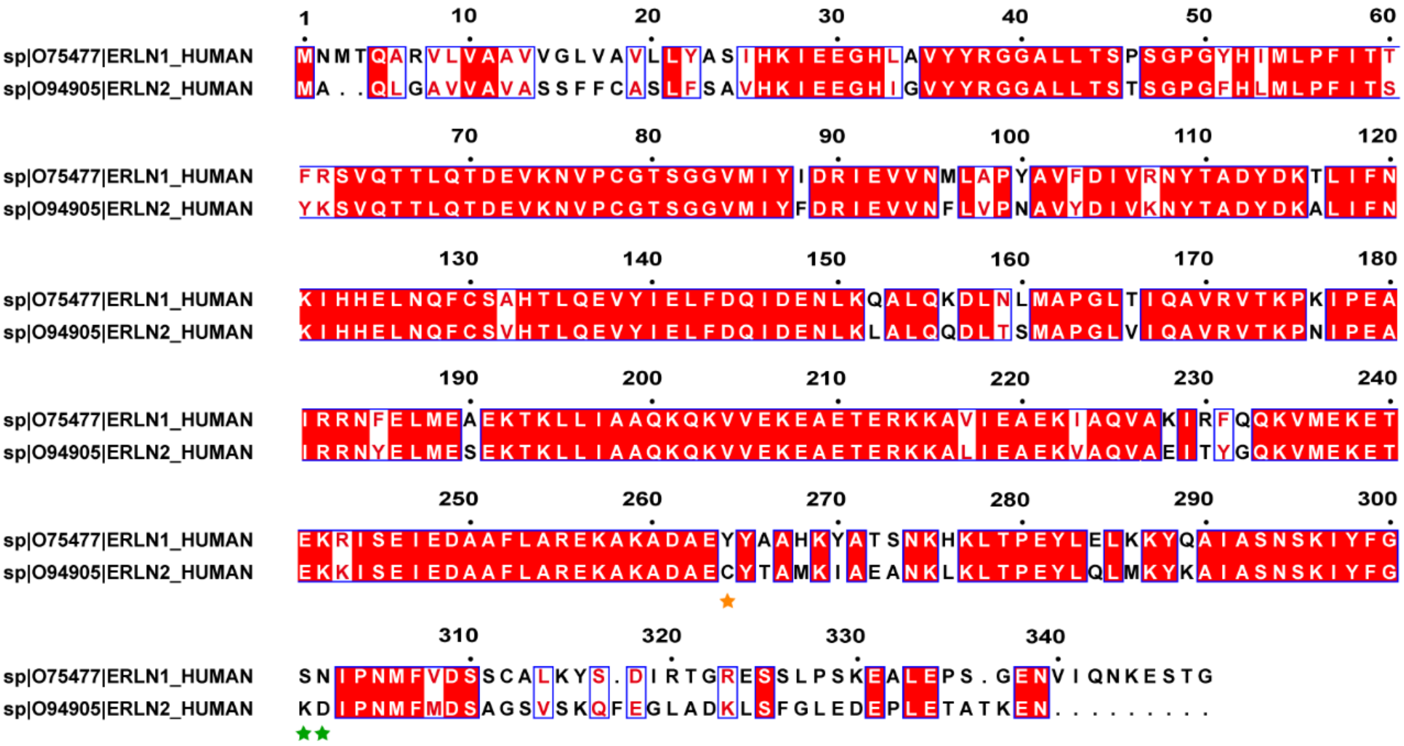

**Figure S1. Sequence alignment of erlin1 and erlin2**

The two proteins share high sequence identity and similarity. The large sequence differences in the N-terminal and C-terminal regions are notable. Non-conserved amino acids in the C-terminal sequence that are important for polymerization are marked with green asterisks. Amino acids of erlin1 and erlin2 with distinguishable density in the cryo-EM map are marked with orange asterisks.

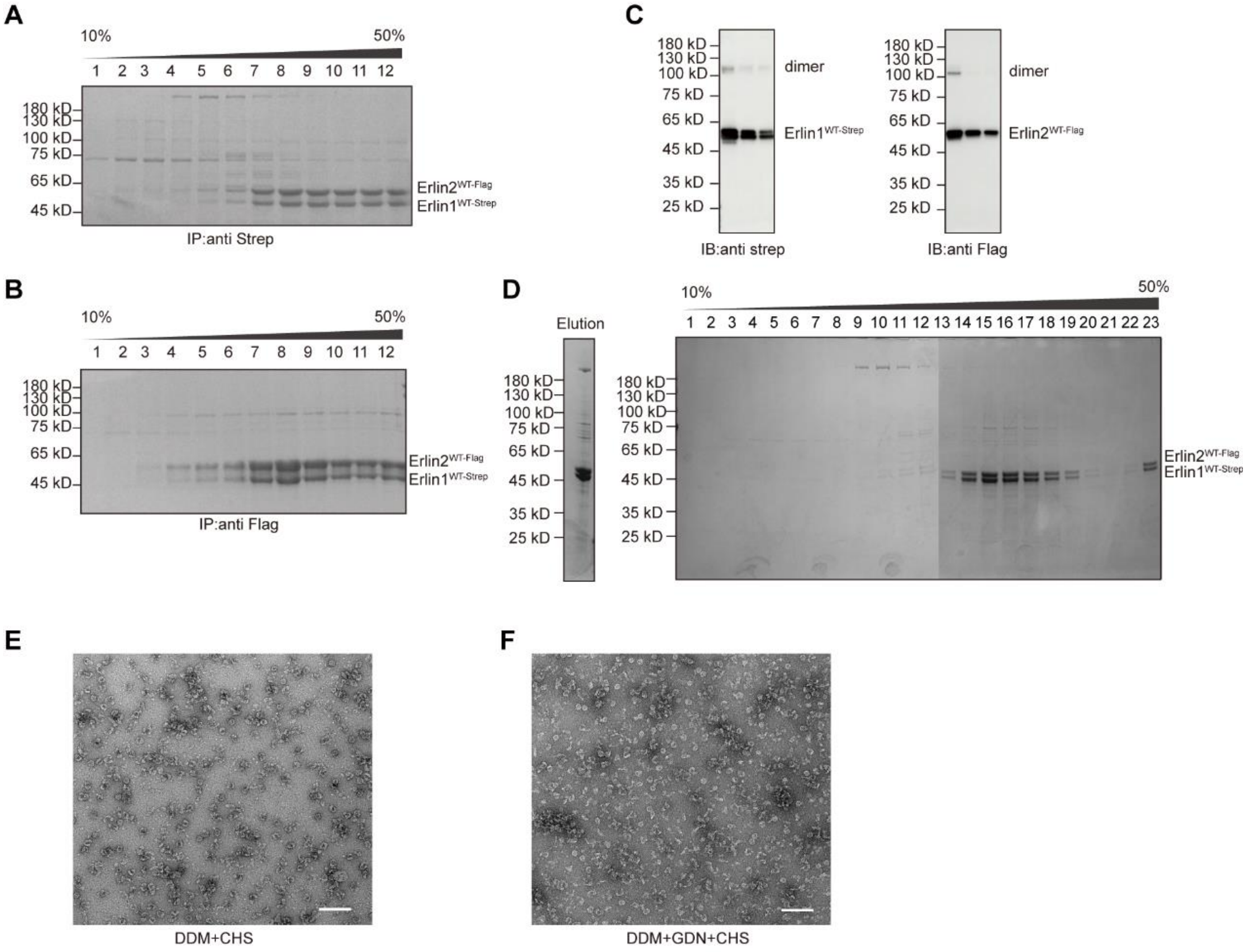

**Figure S2. Purification and detergent condition tests of the erlin1/2 complex**

(A and B) Coomassie Brilliant Blue-stained 10%–50% glycerol gradient fractions of the erlin1/2 complex purified using Strep-tagged erlin1 (A) or Flag-tagged erlin2 (B) under the DDM+GDN solubilization condition.

(C) IB analysis of purified samples detecting erlin1 and erlin2. The three lanes represent different loading amounts.

(D) Eluted sample and 10%–50% glycerol gradient fractions of the erlin1/2 complex purified under the GDN+CHS solubilization condition.

(E and F) nsEM analysis of the erlin1/2 complex purified under various detergent conditions. Protein purification using either DDM+CHS (E) or DDM+GDN+CHS (F) resulted in a high proportion of incomplete particles and greater sample heterogeneity. Scale bar: 200 nm.

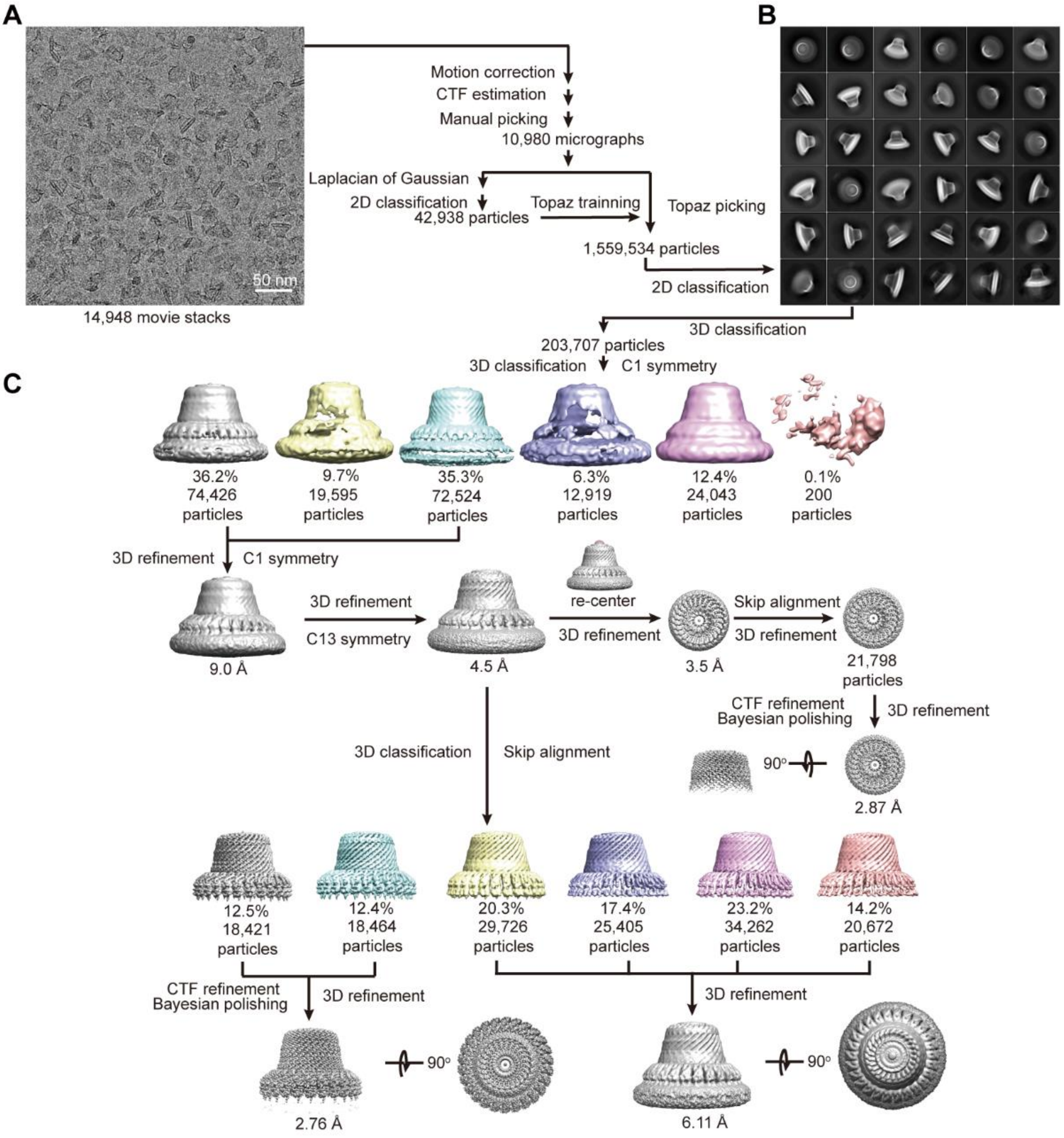

**Figure S3. Data processing of the erlin1/2 complex purified by GDN and CHS**

(A) A representative raw cryo-EM micrograph.

(B) 2D classification of particles binned by 2.

(C) 3D classification and refinement. Approximately 16% of the particles were incomplete. For the complete particles, two final density maps were obtained: one for the expanded cage (6.11 Å) and the other for the contracted cage (2.76 Å). The density of the C-terminal region was improved by mask-based 3D classification (skip alignment), resulting in a map at 2.87 Å resolution.

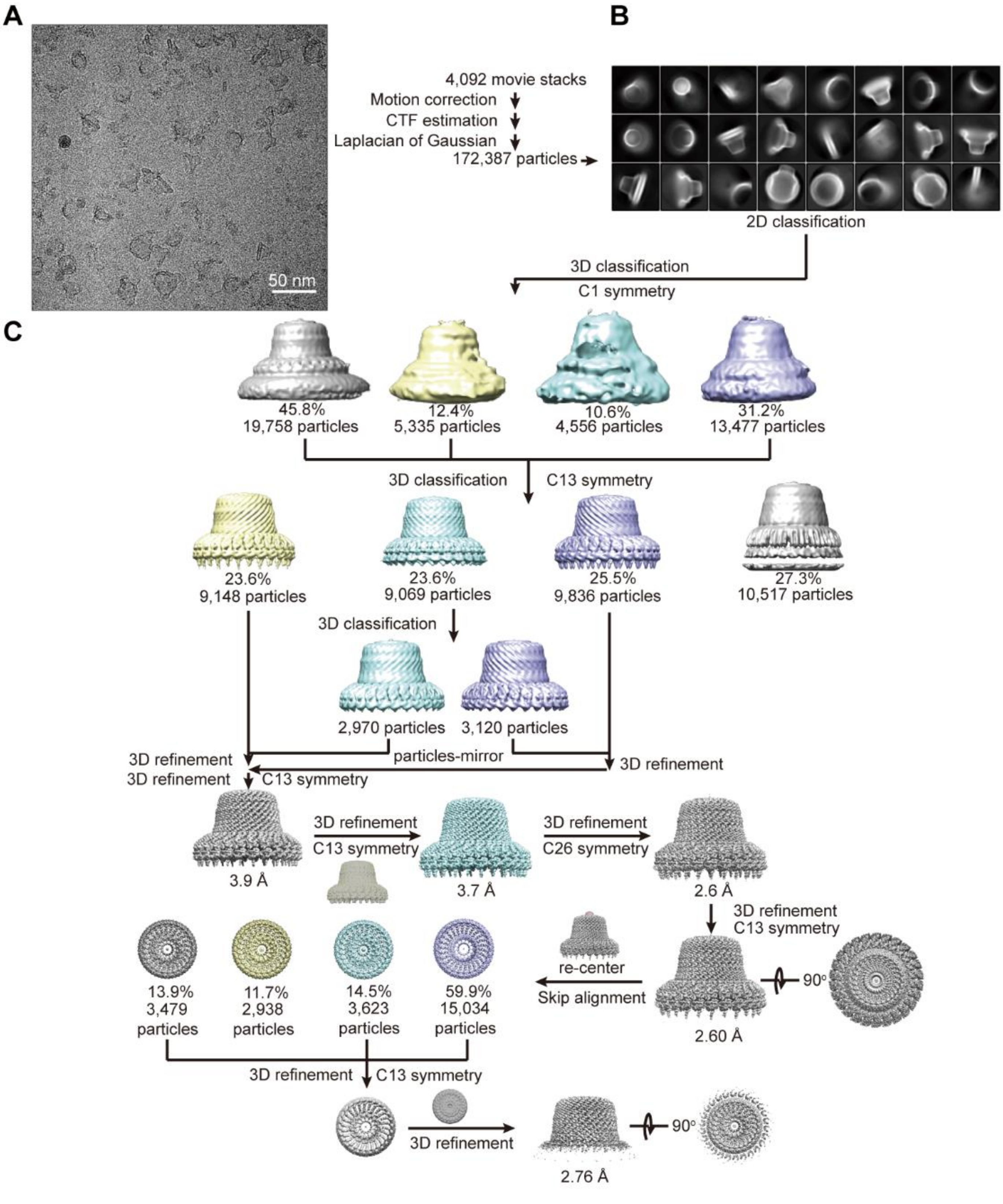

**Figure S4. Data processing of the erlin1/2 complex purified by DDM and GDN**

(A) A representative raw cryo-EM micrograph.

(B) 2D classification of particles binned by 2.

(C) 3D classification and refinement. Approximately 23% of the particles were incomplete. For the complete particles, a final density map was obtained at 2.60 Å resolution. The density of the C-terminal region was improved by mask-based 3D classification (skip alignment), resulting in a map at 2.76 Å resolution.

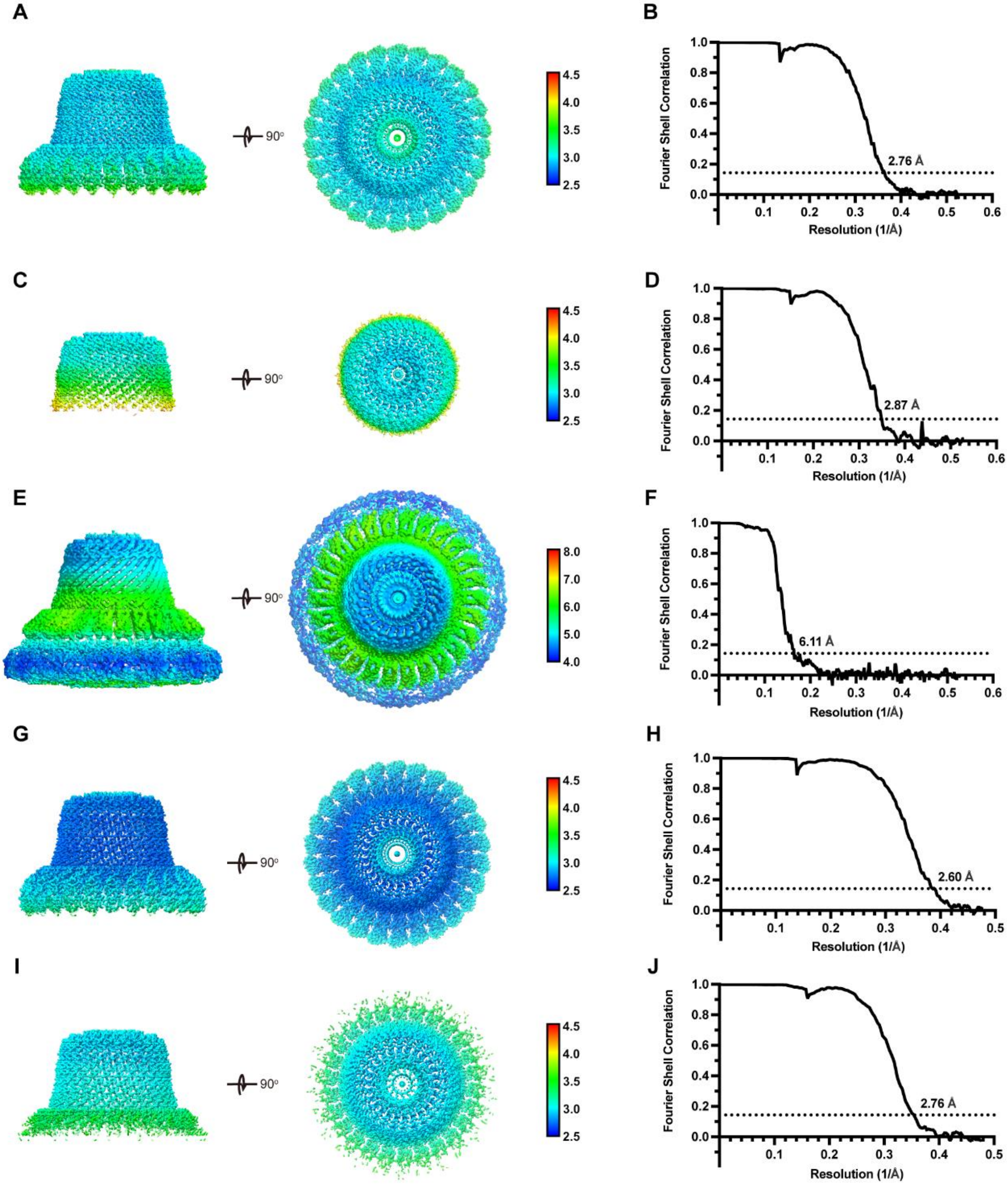

**Figure S5. Resolution estimation of the reconstructed density maps of the erlin1/2 complex**

(A and B) Local resolution estimation (A) and the Fourier shell correlation curve (B) of the erlin1/2 complex in the contracted state obtained under the GDN+CHS solubilization condition.

(C and D) Local resolution estimation (C) and the Fourier shell correlation curve (D) of the C-terminal region of the erlin1/2 complex obtained under the GDN+CHS solubilization condition.

(E and F) Local resolution estimation (E) and the Fourier shell correlation curve (F) of the erlin1/2 complex in the expanded state obtained under the GDN+CHS solubilization condition.

(G and H) Local resolution estimation (G) and the Fourier shell correlation curve (H) of the erlin1/2 complex obtained under the DDM+GDN solubilization condition.

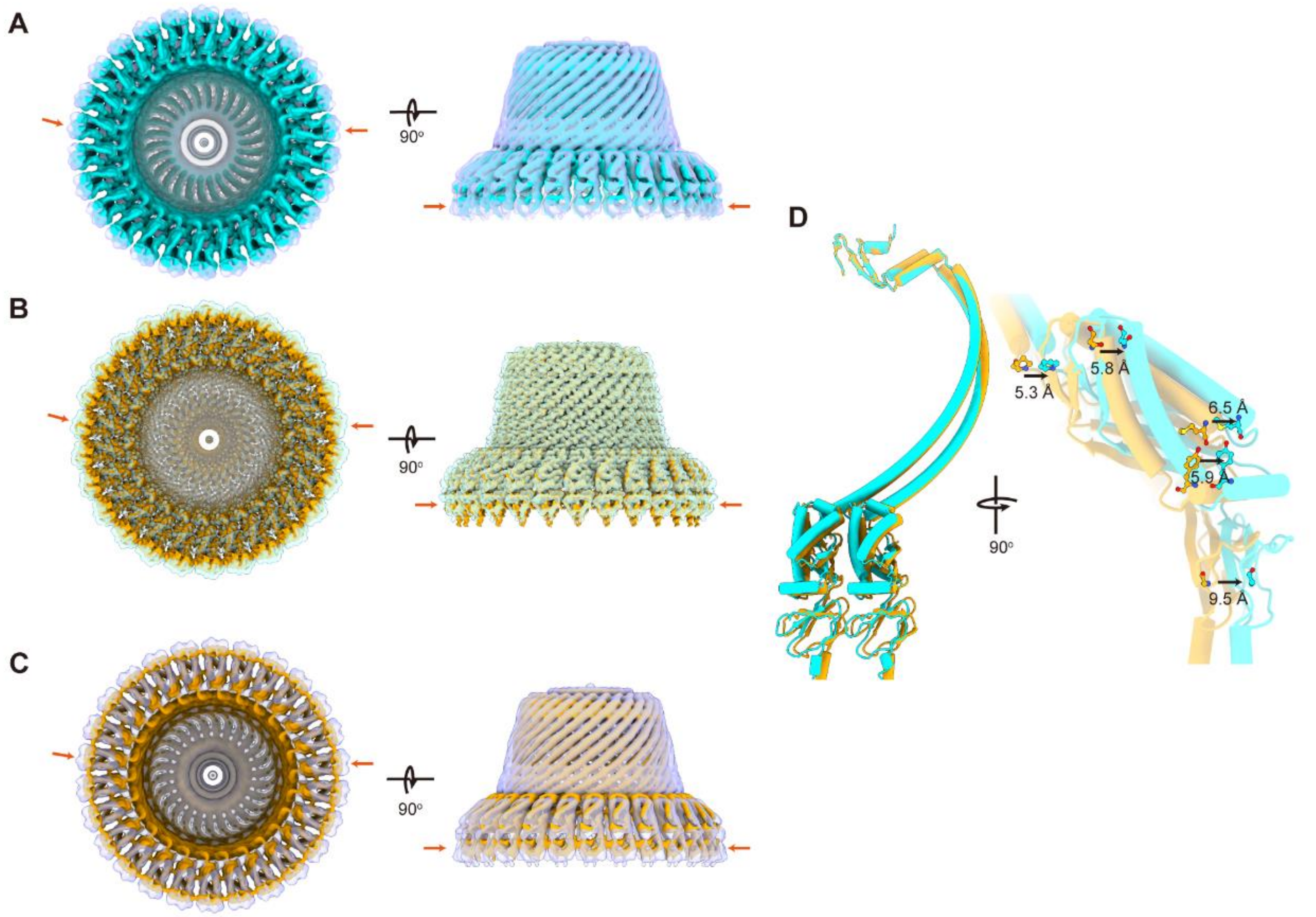

**Figure S6. Comparison of the erlin1/2 structures purified using different detergents**

(A) Bottom (left) and side (right) views of the structural superimposition of the two conformational states of the erlin1/2 cage purified under the GDN+CHS condition. The cyan density map represents the high-resolution map (filtered to 5 Å) of the contracted state, while the semi-transparent density map represents the low-resolution (6 Å) expanded state map.

(B) Bottom (left) and side (right) views of the structural superimposition of the two high-resolution maps of the erlin1/2 complex purified under two conditions. The solid density map was obtained using the DDM+GDN condition and corresponds to the contracted state, while the semi-transparent density map represents the most contracted state from the GDN+CHS condition.

(C) Bottom (left) and side (right) views of the structural superimposition of the expanded state obtained using the GDN+CHS condition (semi-transparent surface representation) and the most contracted state obtained using the DDM+GDN condition (solid surface representation).

(D) Structural comparison between the atomic models derived from the high-resolution maps in (B). The model from the DDM+GDN condition is shown in gold, and the model from the GDN+CHS condition is shown in cyan. When aligned at the C-terminal region, significant displacement is observed at the N-terminal regions. The N-terminal end of the CC1 domain is shifted by ~5 Å, SPFH2 by ~6 Å, and SPFH1 by ~9.5 Å.

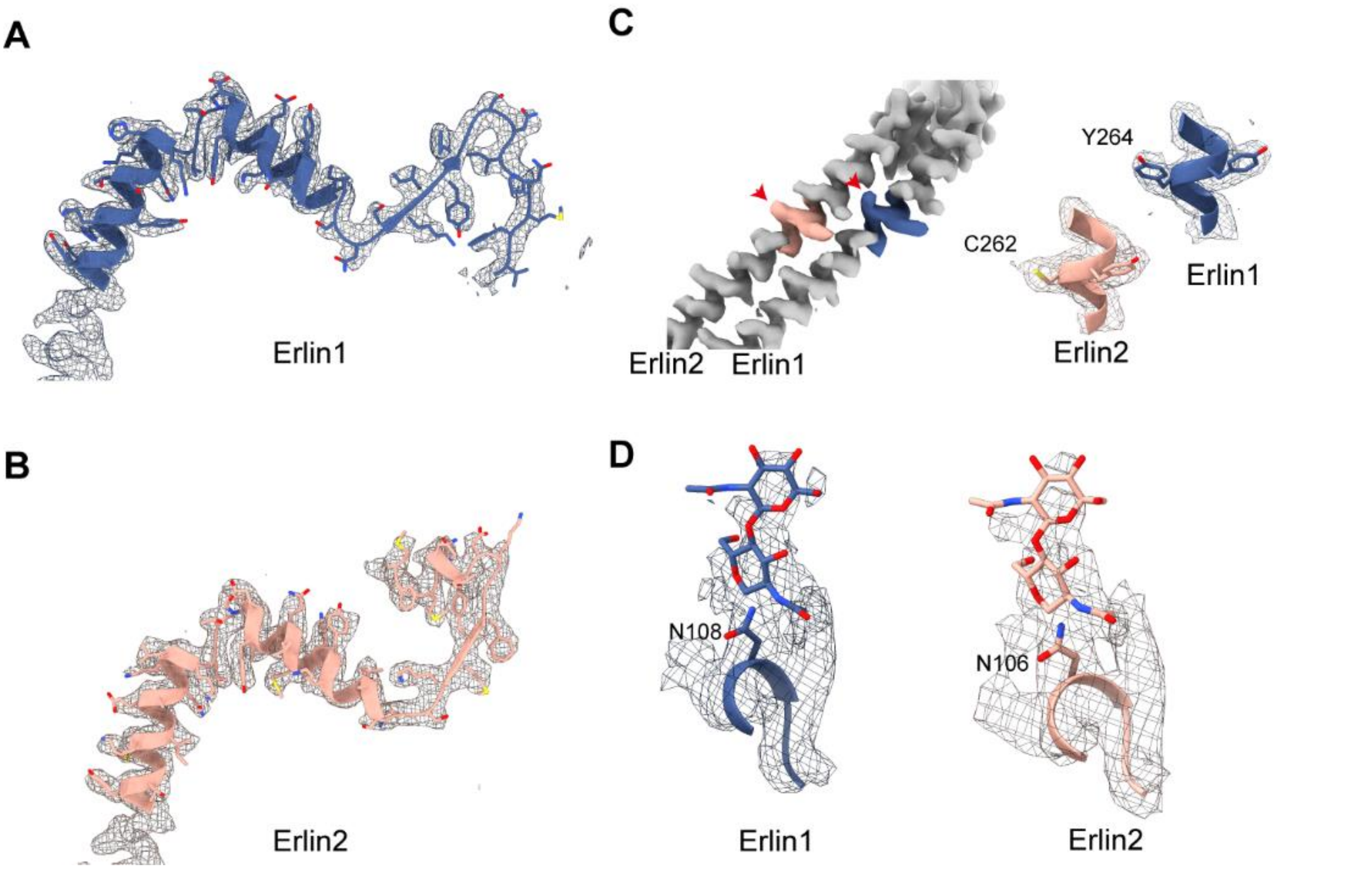

**Figure S7. Local structural differences between erlin1 and erlin2**

(A and B) Cryo-EM density maps and atomic models of the C-terminal regions of erlin1 and erlin2.

(C) Residue Y264 in erlin1 corresponds to C262 in erlin2, showing discernible side-chain density differences.

(D) Glycosylation sites of erlin1 and erlin2. N108 in erlin1 and N106 in erlin2 are predicted to be N-linked glycosylation sites, with the observed extra density accommodating one or two sugar moieties.

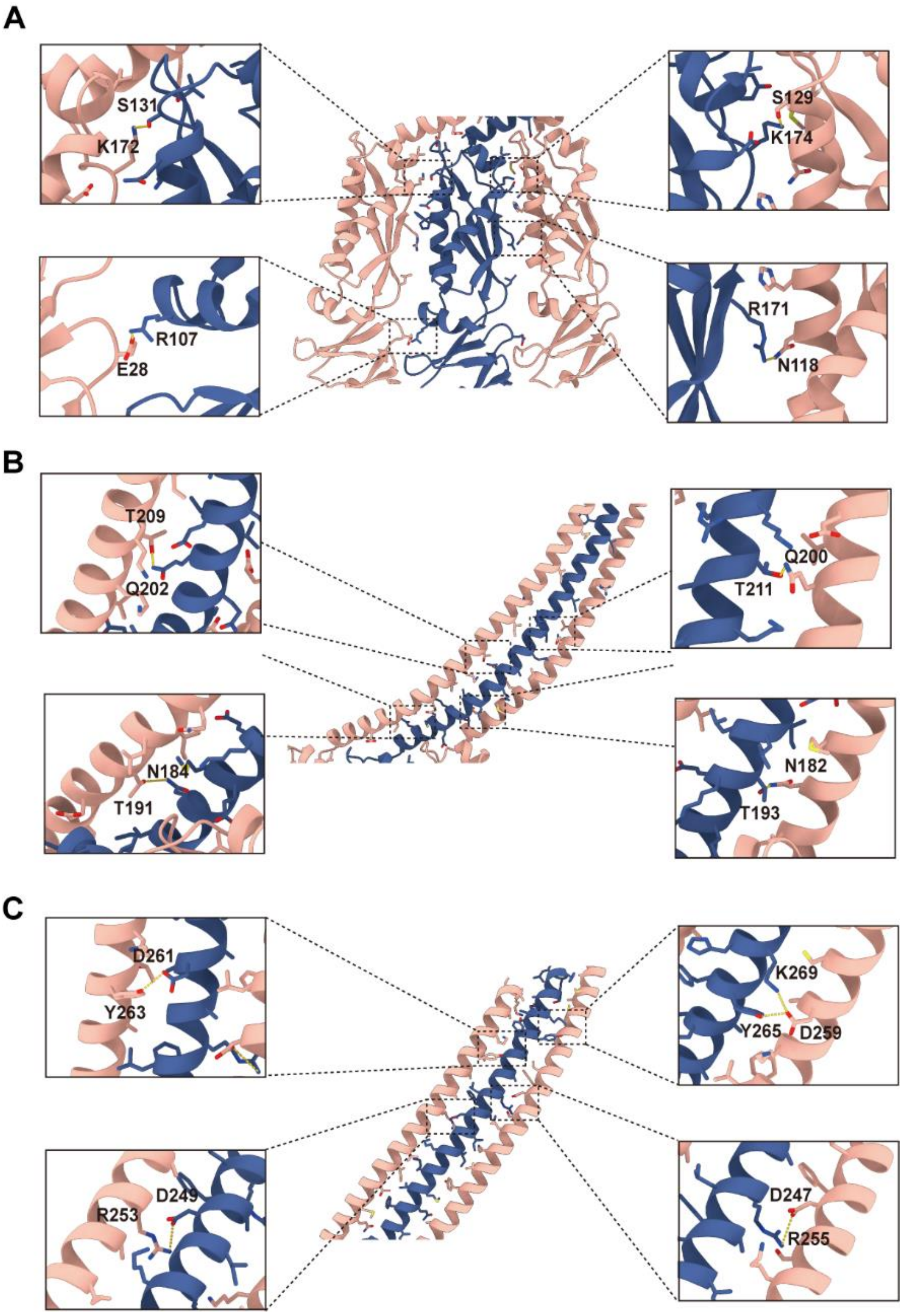

**Figure S8. Interaction between erlin1 and erlin2 in the SPFH and CC1 domains**
(A) Interaction between the SPFH domains of erlin1 and erlin2.
(B and C) Interaction between the CC1 domains of erlin1 and erlin2.

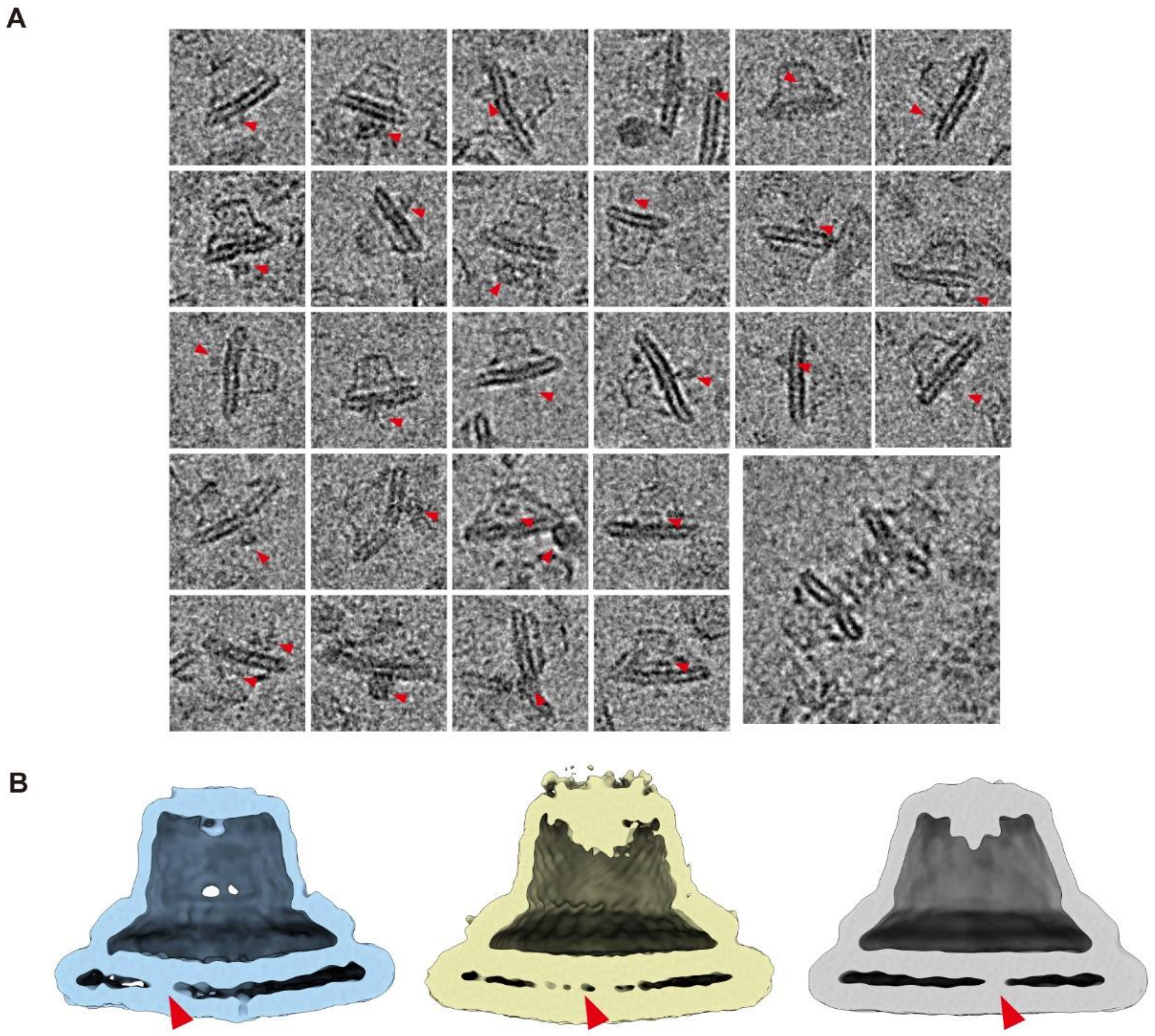

**Figure S9. Presence of additional density in the erlin1/2 complex**

(A) Representative raw cryo-EM particles showing that additional protein densities were observed both inside and outside the cage.

(B) Cross-section views of the reconstructed density maps of the erlin1/2 complex. 3D classification with C1 symmetry revealed extra transmembrane density within the membrane region of the cage.

**Supplementary Table 1**

Guide RNA sequences for erlin1 and erlin2 knockout

| | Guide sequence (5' - 3') |
|---|---|
| *Lacz* sgRNA | TGCGAATACGCCCACGCGAT |
| mouse *ERLIN1* sgRNA1 | TGTGGATGGAGGCGTACAGG |
| mouse *ERLIN1* sgRNA2 | TATGATAGCCTGGTCCACTG |
| mouse *ERLIN2* sgRNA1 | GCTGTGCACAAGATAGAAGA |
| mouse *ERLIN2* sgRNA2 | GCCCTGCTGACCTCCACCAG |
| human *ERLIN1* sgRNA1 | TATGATAGCCTGGTCCACTG |
| human *ERLIN1* sgRNA2 | AGAACTGGTTCAGCTCATGG |
| human *ERLIN2* sgRNA1 | GTAGGTGATCTCAGCCACCT |
| human *ERLIN2* sgRNA2 | AGAACTGGTTCAGTTCGTGG |

**Supplementary Table 2**

Oligonucleotide Sequences for Quantitative PCR (qPCR)

| | Forward (5’ - 3’) | Reverse (5’ - 3’) |
|---|---|---|
| mouse β-Actin | GGCTGTATTCCCCTCCATCG | CCAGTTGGTAACAATGCCATGT |
| human β-Actin | TTTCTGTCACTCTTCTCTTAGGT | AGGTCTTTACGGATGTCAAGG |
| MHV-N | CAGATCCTTGATGATGGCGTAGT | AGAGTGTCCTATCCCGACTTTCTC |

**Supplementary Table 3**

Cryo-EM data collection, refinement and validation statistics of the erlin1/2 complex

| | The erlin1/2 complex (DDM+GDN) (EMD-XXXX) (PDB: YYYY) | The erlin1/2 complex (GDN+CHS) (EMD-XXXX) (PDB: YYYY) |
|---|---|---|
| **Data collection and processing** | | |
| Electron microscope | Titan Krios | Krios G4 |
| Electron detector | Gatan K2 | Falcon 4 |
| Magnification | 130,000× | 130,000× |
| Voltage (kV) | 300 | 300 |
| Electron exposure ($e^{-}/Å^{2}$) | 64 | 54.12 |
| Defocus range (μm) | -1.2 to -1.8 | -1.2 to -1.8 |
| Pixel size (Å) | 1.052 | 0.95 |
| Micrographs | 4,209 | 14,948 |
| Symmetry imposed | C13 | C13 |
| Initial particle images | 196K | 1559K |
| Final particle images (total) | 25K | 146K |
| Map resolution (total) (Å) | 2.60 | 2.76 |
| FSC threshold | 0.143 | 0.143 |
| **Refinement** | | |
| Model composition | | |
| Non-hydrogen | 63388 | 62374 |
| atoms | 7904 | 7748 |
| Protein residues | 52 | 52 |
| Ligands | | |
| R.m.s. deviations | | |
| Bond lengths (Å) | 0.003 | 0.003 |
| Bond angles (°) | 0.579 | 0.622 |
| Validation | | |
| MolProbity score | 1.75 | 1.66 |
| Clashscore | 6.37 | 4.45 |
| Rotamer outliers (%) | 3.28 | 3.18 |
| Ramachandran plot | | |
| Favored (%) | 97.99 | 97.74 |
| Allowed (%) | 1.88 | 2.25 |
| Outliers (%) | 0.13 | 0.01 |
